# A visual atlas of meiotic protein dynamics in living fission yeast

**DOI:** 10.1101/2020.06.15.153338

**Authors:** Wilber Escorcia, Vishnu P. Tripathi, Ji-Ping Yuan, Susan L. Forsburg

**Affiliations:** University of Southern California, Molecular and Computational Biology Program; University of Southern California, Leonard Davis School of Gerontology; Xavier University, Department of Biology

**Keywords:** Meiosis, Histone, Cohesin, Microtubule, Spindle Assembly Checkpoints, PCNA

## Abstract

Meiosis is a carefully choreographed dynamic process that re-purposes proteins from somatic/vegetative cell division, as well as meiosis-specific factors, to carry out the differentiation and recombination pathway common to sexually reproducing eukaryotes. Studies of individual proteins from a variety of different experimental protocols can make it difficult to compare details between them. Using a consistent protocol in otherwise wild type fission yeast cells, this report provides an atlas of dynamic protein behavior of representative proteins at different stages during normal zygotic meiosis in fission yeast. This establishes common landmarks to facilitate comparison of different proteins and shows that initiation of S phase likely occurs prior to nuclear fusion/karyogamy.

**Summary:** Meiosis is an important process for sexually reproducing organisms. Unique dynamics of recombination and chromosome segregation are required for this differentiation process. Fission yeast is an excellent model to study meiotic progression and chromosome dynamics. Historically, different methodologies have been used to examine protein dynamics in fixed or live cells, which makes comparisons more difficult. In this report, we use fluorescently tagged proteins and live-cell microscopy under uniform conditions to compare meiotic signposts that define dynamic behavior of proteins during meiotic DNA synthesis, nuclear fusion, chromosome alignment, genetic recombination, metaphase, and meiosis. This establishes a reference atlas of protein behavior during meiotic differentiation.

## Introduction

Meiosis is a conserved cell differentiation pathway found in sexually reproducing eukaryotes. It is characterized by one round of DNA synthesis, followed by two nuclear divisions, resulting in haploid gametes (reviewed in (Zickler and Kleckner, 1999); (Cavalier-smith, 2002; Hochwagen, 2008; Ohkura, 2015)). In most organisms, the divisions are preceded by programmed double strand breaks (DSB) followed by homologous recombination (HR) that facilitates separation of the homologous chromosomes during reductional meiosis I division (reviewed in (Gray and Cohen, 2016)). The equational meiosis II division separates sister chromatids and more closely resembles mitosis in somatic or vegetative cells but is unusual in that it does not follow a period of DNA synthesis (reviewed in (Ohkura, 2015)).

A variety of meiosis-specific genes are recruited to facilitate these specialized events. These include additional recombination proteins, components of the synaptonemal complex or linear elements between chromosome homologues, proteins that contribute to kinetochore mono-orientation, and meiosis-only cohesin proteins that link sister chromatids together. These specialists modify the basic architecture of cell division particularly to facilitate the meiosis-specific chromosome recombination and the MI division mechanisms (Hunter, 2015; Lam and Keeney, 2015; Zickler and Kleckner, 2015). The yeast *Schizosaccharomyces pombe* offers an excellent model system to study the process of meiosis. Normally haploid cells can be induced to conjugate with cells of the opposite mating type in conditions of nutrient limitation, forming transient zygotes. The initial landmark event following karyogamy is the formation of a telomere-led chromosome bouquet, which migrates rapidly back and forth in the length of the cell in a dynein-led movement called horse-tailing (HT) (Ding et al., 1998; Yamamoto et al., 1999). Meiotic S (meiS) phase and recombination occur during horse-tailing, which is followed by the MI and MII divisions (reviewed in (Davis and Smith, 2001; Mizuguchi et al., 2015; Yamashita et al., 2017). Genetic tools have been deployed to identify and characterize proteins that function throughout these different phases.

The advent of green fluorescent protein (GFP) and related colorful fluorescent tags has allowed the analysis of protein dynamics in living cells and these have been widely used in studies of fission yeast meiosis. However, variation in protocols and conditions make it challenging to compare the dynamics of proteins in the various stages of meiosis between different publications. In particular, many studies employ a temperature sensitive *pat1* mutant to drive a synchronous meiosis, even from haploids, but *pat1* disrupts some aspects of meiosis (e.g., (Bähler et al., 1991; Iino and Yamamoto, 1985a; Yamamoto and Hiraoka, 2003). In this report, we examine protein dynamics in meiosis using consistent strains, growth conditions and a standardized imaging protocol (Escorcia et al., 2019; Green et al., 2015). This allows us to generate an atlas of meiotic protein behavior that will be a useful reference to many investigators studying this dynamic process.

## Results and Discussion

**Rationale** We used a uniform growth strategy to induce mating and meiosis between cells of opposite mating types, followed by imaging of cells on agar pads (see Methods (Escorcia and Forsburg, 2017; Escorcia et al., 2019)). Importantly, aside from the tagged genes, the strains are normal haploids that were induced to mate, followed directly by zygotic meiosis. Reasonable synchrony was achieved solely by using appropriate growth conditions, and we did not employ any mutations such as *pat1-ts* which can induce meiotic abnormalities (e.g., (Bähler et al., 1991; Yamamoto and Hiraoka, 2003)). Thus, our strategy recapitulates as much as possible a “normal” fission yeast meiosis.

### Nuclear dynamics establish a reference

Hht1 and Hhf1 are the H3-H4 histone pairs that package DNA in fission yeast (Choe et al., 1985; Takayama and Takahashi, 2007). Because of its role in nuclear organization, Hht1 is used with fluorescent tags to mark the nuclear mass during mitotic and meiotic processes (Escorcia and Forsburg, 2017; Sabatinos et al., 2015; Tomita et al., 2013). To establish an initial reference for meiotic progression, we employed cells carrying homozygous Hht1-mRFP and followed meiotic progression using live-cell imaging. We began our observations prior to karyogamy, during Pre-Fusion (PF), and followed through karyogamy/nuclear fusion (FS), horse-tailing (HT), metaphase I (MTI), meiosis I and all the way up-to meiosis II (MII).

To set a common landmark for timing comparisons, we used the clearly recognizable characteristics of metaphase I (MTI) as a reference time = 0 (t0) minute (MTI: 0′). Events occurring prior to MTI are represented by negative timing numbers (e.g. −10′, −20′ etc.) and events occurring post MTI are by positive numbers (e.g. 10′, 20′ etc.).

In our initial examination of histone dynamics, we observed a bright, pan-nuclear Hht1-mRFP signal at prefusion (PF) (PF: −210′) with no signs of vigorous nuclear movements (**Fig. 1A Fig. S1A)**. At karyogamy/nuclear fusion (FS), we observed a characteristic nuclear bridge structure, mixing of parental nuclei, and doubling of nucleus size in the zygote relative to pre-fusion cells (FS: −190′). The period of nuclear fusion lasts for 14 minutes (mean timing= 14.1′± 2.6′) before the nuclear mass undergoes repeated oscillations for approximately 215 minutes (215.5′± 15.8′), representing horse-tailing (**Fig. 1B**). When horse-tailing concludes, vigorous oscillation ceases, and the nuclear size starts to contract, with moderate to low nuclear movement (HT to MTI: −30′ to −10′) (**Fig. 1A, Fig. S1 B-C**). This transition phase of nuclear reconfiguration lasts for approximately 20 minutes (23.2′± 1.7′) followed by 8 minutes (8.4′± 1.8′) of metaphase I (MTI) where maximal nuclear compaction occurs and Hht1-mRFP signal intensifies (HT-MTI: −20′ to 0′) (**Fig. 1 A-C**).

**Figure 1.**
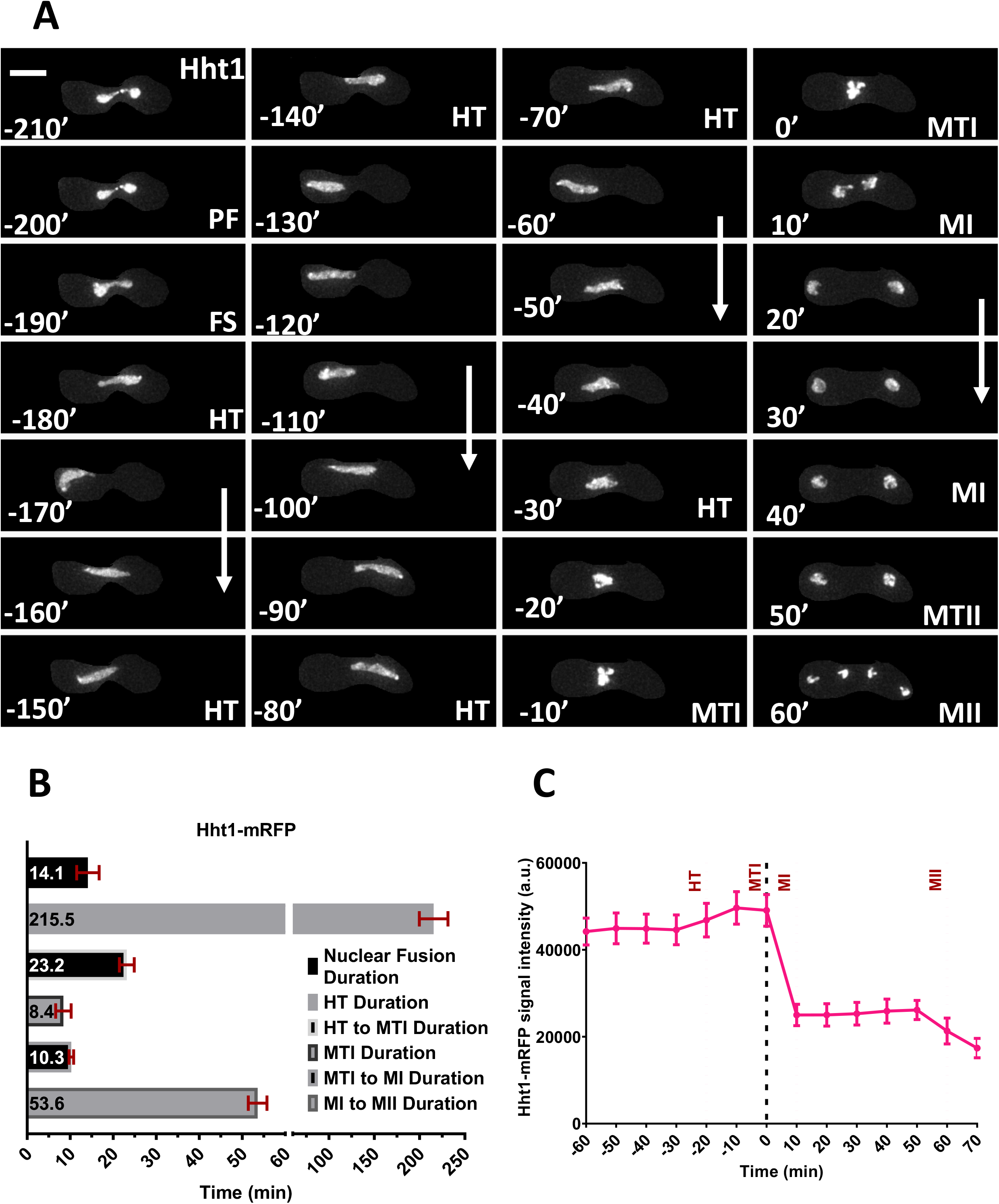
Nuclear dynamics of Hht1 defines signposts for meiotic events. (A) Livecell image of a heterothallic unperturbed zygotic meiotic cell carrying fluorescently tagged histone 3 (Hht1-mRFP) (FY5608 × FY5609). Time lapse images were captured every 10 minutes for 8 hours and selected frames are shown. More than 35 cells from at least two independent biological replicates movies were analyzed and representative image presented. Panel showing hht1-mRFP dynamics during meiotic cycle encompassing different meiotic signposts as pre-fusion (PF: −210′ to −200′), karyogamy (FS: −190′), horse-tailing (HT: −180′ to −20′), metaphase I (MTI: −10′ to −0′), meiosis I (MI: 10′ to 40′), metaphase II (MTII: 50′) and meiosis II (MII: 60′). Numbers are representing timing in minutes and the scale bars are equal to 5 μm. Metaphase I, just prior to meiosis I is considered as “t0” and any event occurring prior to that are represented by negative timing numbers while post metaphase I events are marked by positive numbers (B) The durations of Hht1-mRFP dynamics in different meiotic event. (C) Quantitation of Hht1-mRFP fluorescence intensity. Average (mean) values of timing and fluorescence intensities are presented in (B) and (C) respectively. Error bars represent 95% class intervals.

To better represent the changes in hht1-mRFP signal intensities along with nuclear dynamics, we used the “Thermal” LUT feature of FIJI. In this presentation, the yellow and red color signals of hht1-mRFP represent higher intensities due to higher compaction of the nucleus (**Fig. S1A**). The nucleus in meiosis I (MI) splits into two nuclear masses of roughly equal size (MI: 10′) (**Fig. 1 A, Fig S1B**) that contain segregated sets of homologous chromosomes (Hauf et al., 2007; Hirose et al., 2011; Yokobayashi and Watanabe, 2005). The transition of metaphase I to meiosis I lasts for approximately 10 minutes (10.3′± 0.5′), whereas the average duration of meiosis I to meiosis II is approximately 50 minutes (53.2′± 2.2′) (**Fig. 1A-B**). During metaphase II (MTII), homologs continue to move toward opposite ends of the zygote. Final separation of the two nuclear masses into four daughter nuclei marks the completion of meiosis II (MII: 60′) (**Fig. 1 A**). At this stage, sister chromatid sets completely segregate and are readied for packaging into spores in the gamete-maturation phase that follows meiosis II (Hauf et al., 2007; Hirose et al., 2011; Yokobayashi and Watanabe, 2005).

Our observations of H3 histone (Hht1-mRFP) dynamics generally agree with those of other live-cell studies (Ding et al., 2006; Escorcia and Forsburg, 2017; Fennell et al., 2015; Klutstein et al., 2015; Tomita et al., 2013; Yamamoto et al., 2019; Yang et al., 2015). The timing from nuclear fusion to meiosis II takes approximately 4 ½ hours, which is in line with previous reports (Ding et al., 2006; Moiseeva et al., 2017; Tomita et al., 2013). These histone dynamics established a baseline that we used to compare different strains and markers.

We observed that predictable changes in nuclear mass correlate with changes that define functional shifts in different markers of meiotic progression. As expected, there is a correlation between signal intensity changes and changes in nuclear size. Significant increase in Hht1-mRFP fluorescence intensity are observed during fusion and in each of the two metaphase steps preceding the meiotic divisions. This link is consistent with reports showing elevated histone content during DNA synthesis and increased nuclear condensation in metaphase (Chikashige et al., 2004; Choe et al., 1985; Ding et al., 2004; Takayama and Takahashi, 2007; Yokobayashi et al., 2003).

### S phase markers PCNA and Tos4

PCNA (SpPcn1) plays major roles in DNA replication, repair and translesion synthesis (revived in (Boehm et al., 2016). As a processivity factor, it interacts with both DNA Polymerase *δ* and *ε* (Arroyo et al., 1996; reviewed in (Choe and Moldovan, 2017)). To study the nuclear dynamics and timings of meiotic S phase, we used a heterozygous cross between cells containing EGFP-Pcn1 (Meister et al., 2007) and hht1-mRFP. For the S phase markers, instead of metaphase I, we used the initial nuclear fusion stage (FS) to represent the t0 minute (FS: 0′), to resolve earlier events. Events prior to karyogamy or prefusion have negative timing (PF: −10′, −20′) and post fusion stage such as horse-tailing stage has positive timing numbers (HT: 10′, 20′ etc.)

During PF stage (PF: −10′), we observed bright and discrete EGFP-Pcn1 foci, similar to those observed in vegetative S phase (Meister et al, 2007). These were distributed across both nuclei and persisted for about 10 minutes (11.1 ± 1.1′) (**Fig. 2A, B**). Notably, the histone signal is observed in just one parent, further indicating that fusion has not occurred. During karyogamy (FS: 0′), the foci remained distributed across the nucleus and the signal intensity increased (**Fig. 2C**). As cells moved from karyogamy to the horse-tailing stage, EGFP-Pcn1 signal remains distributed across the nucleus (HT: 10′) but localizes in one part, which may indicate late replication activity in certain parts, possibly the nucleolus (HT: 10′ to 30′) (**Fig. 2A)**. We observed an average of about 49 minutes (48.9′± 8.1′) from karyogamy before the EGFP-Pcn1 signal is lost. Notably, horse-tailing continues for another 160-180 minutes but Pcn1 signal remains diffused, which we infer indicates the conclusion of S phase.

**Figure 2.**
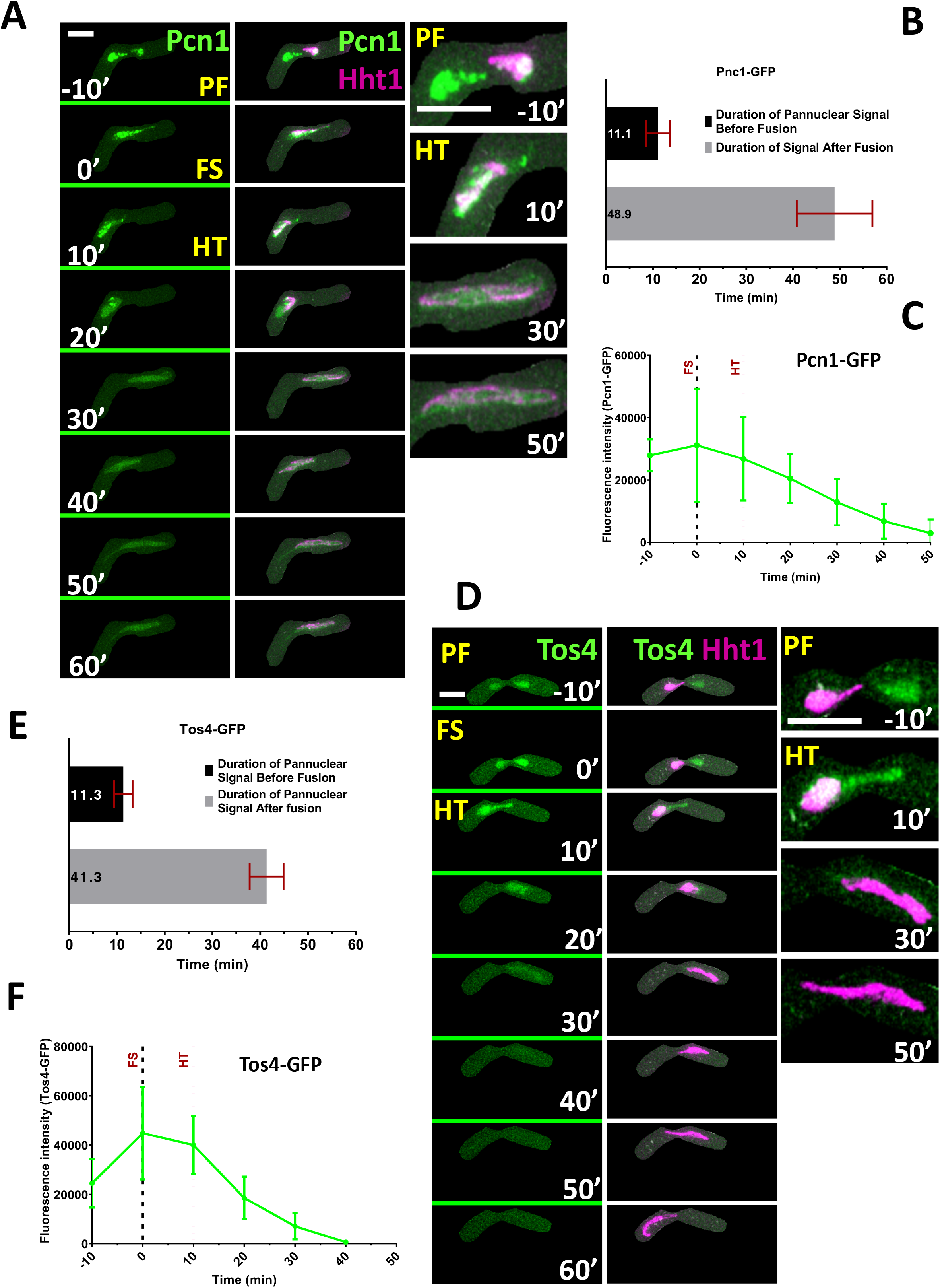
Nuclear dynamics of Pcn1 and Tos4 during meiotic S phase. (A) Live-cell images of zygotic meiotic cell carrying EGFP tagged PCNA (EGFP-Pcn1) and mRFP tagged Hht1 (3050 × 5615). Time lapse images were captured every 10 minutes for 8 hours and selected frames are shown. Panel showing dynamics of EGFP-Pcn1 (column 1) and EGFP-Pcn1, Hht1-mRFP merged (column 2) during early meiotic events encompassing Pre-fusion (PF), Karyogamy (FS) and early horse-tailing (HT) stage. Column 3 is showing closeups of selected time points from column 2. Exclusively for S phase markers, the “t0” is changed from metaphase I (MTI) to fusion (FS) stage. Events prior to FS are represented by negative numbers and post fusion events are by positive timing numbers. Numbers represent timing in minutes and scale bars equal 5 μm. (B) Durations of EGFP-Pcn1 foci signal disappearance. (C) Quantitation of EGFP-Pcn1 fluorescence intensity. (D) Time lapse images of a zygotic meiotic cells carrying Tos4-GFP and Hht1-mRFP (8848 × 5615). Panel showing dynamics of Tos4-GFP (column 1), Tos4-GFP and hht1-mRFP merged (column 2) during early meiotic events encompassing Pre-fusion (PF), Karyogamy (FS) and early horse-tailing (HT) stage. Column 3 is a closeups of selected time points from column 2. Numbers represent timing in minutes and scale bars equal to 5 μm. (E) Durations of Tos4-GFP signal disappearance. (F) Quantitation of Tos4-GFP fluorescence intensity. For estimating timing and signal intensity, n=9 (PCNA) and n=15 (Tos4) cells from at least two independent biological replicate movies were analyzed. Data in (B), (C), (E) and (F) are representing mean values. Error bars represent 95% class intervals.

Previously, in a *pat1*-induced meiosis, it was reported that DNA replication occurs at the beginning of the horsetail stage (Chikashige et al., 2004). However, a previous study using fluorescently tagged Pcn1 in homothallic *h90,* is consistent with our study using heterothallic *h^-^* and *h^+^*, suggesting early S phase begins prior to karyogamy (Ruan et al., 2015). We speculate that initiation of premeiotic S phase is likely coordinated by trans-acting transcription factors that form after cell fusion but prior to nuclear fusion (e.g.,(Vještica et al., 2018)).

We examined another S-phase marker, Tos4, which is a transcription factor that accumulates in the nucleus during early S-phase (Bastos de Oliveira FM et al., 2012; Kiang et al., 2009; Kim et al., 2020; Shen and Forsburg, 2019). It contains a fork-head associated (FHA) domain and is regulated by the G1/S transcription factors SBF (Swi4-Swi6 cell cycle box binding factor) (Horak et al., 2002; Huang and Elledge, 2000). Here again we used a heterozygous cross, with one parent including fluorescently tagged Tos4 (Tos4-GFP) and the other Hht1 (Hht1-mRFP), to examine its behavior in meiosis. Like EGFP-Pcn1we observed bright foci distributed throughout the nucleus prior to nuclear fusion (PF: −10′), which lasted for about 11 minutes (11.3′± 1.9′) (**Fig. 2D, E**). As cells moved to karyogamy, the signal intensity increased and reached to maximum (FS: 0′) (**Fig. 2F**). The Tos4-GFP signal gradually disappears as cells spend about 25-30 minutes in horse-tailing (HT: 30′) (**Fig. 2D, F**). The duration of Tos4 signal from fusion to early horse-tailing is observed for about 40 minutes (41.3′± 3.5′) (**Fig. 2E**). The initial dynamics and timing of Tos4-GFP accumulation in the nucleus is similar to those of EGFP-Pcn1, but without late focus formation during the later phase of HT with Pcn1 (Fig. 2A: HT: 20′). This suggests that Tos4-GFP is an early meiotic S phase marker with similar dynamics to vegetative S phase (Kim et al., 2020).

The consistent nuclear dynamics and timing of the two meiotic S phase markers, Tos4 and Pcn1, indicate that the activation of meiotic DNA replication starts prior to nuclear fusion. Indeed, given the approximately 25 minutes maturation time of GFP (Heim et al., 1995), it is likely to initiate even earlier than our observation.

### Cohesin and pairing proteins

Rec 8 is the alpha-kleisin subunit of the meiotic cohesin complex in fission yeast (Ding et al., 2006; Ding et al., 2016; Watanabe and Nurse, 1999; Watanabe et al., 2001). Following meiotic DNA synthesis, it keeps sister chromatids cohered at the centromere until just before meiosis II, when it is degraded to allow for sister chromatid separation (Ishiguro et al., 2010; Kitajima et al., 2004; Kitajima et al., 2006; Riedel et al., 2006). We examined meiotic progression in strains in a homozygous cross with Rec8-GFP and Hht1-mRFP (**Fig. 3**).

**Figure 3.**
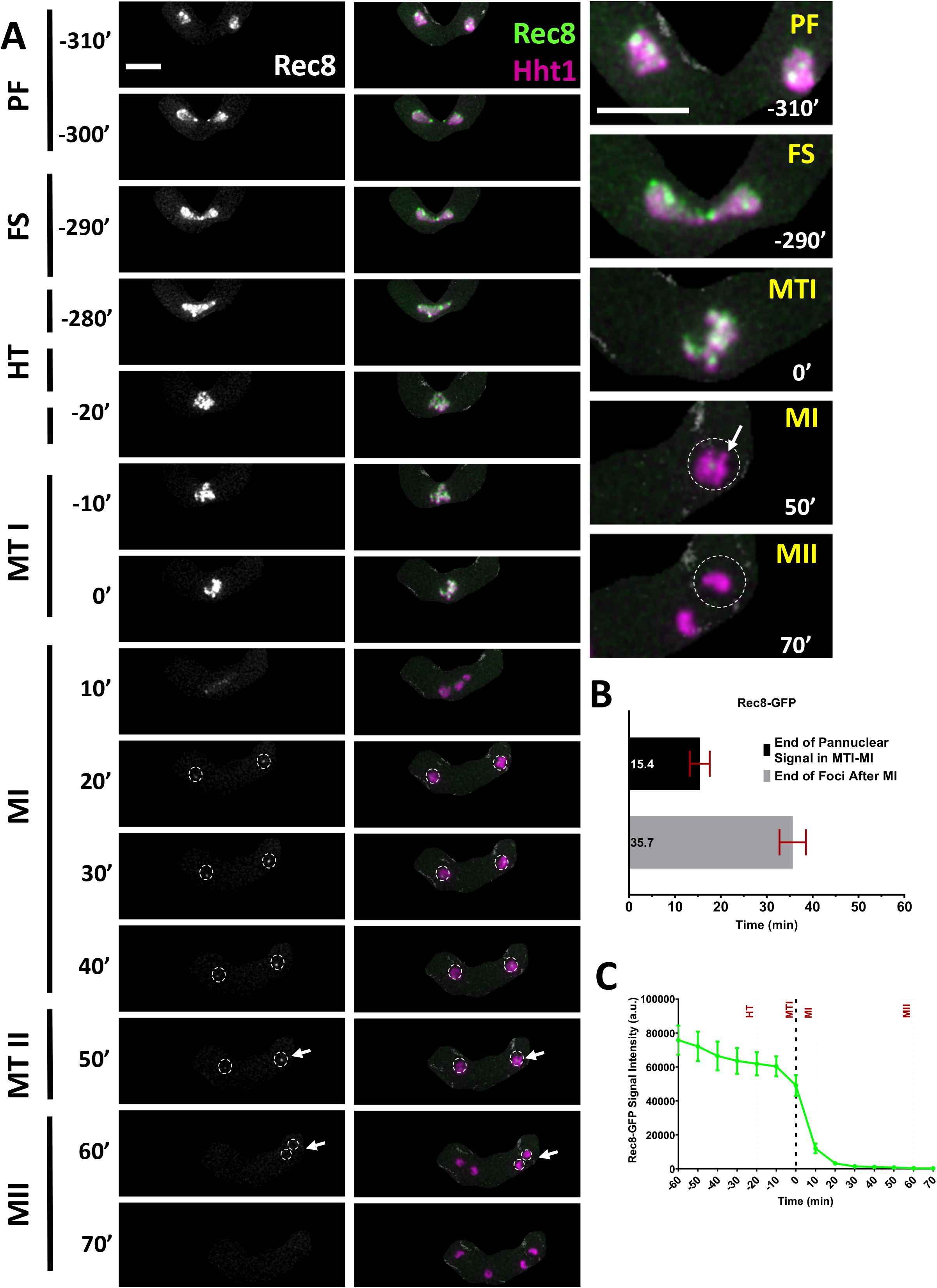
Rec8 cohesin dynamics. (A) Time-lapse images of meiotic cells carrying homozygous Rec8-GFP and Hht1-mRFP (7748 × 7840) were captured every 10 minutes for 8 hours and selected frames are shown. Panel showing cohesin dynamics, (column 1:Rec8-GFP) and (column 2: merged Rec8-GFP Hht1-mRFP), during different meiotic events covering pre-fusion (PF: −310′ to −300′), karyogamy (FS: −290′), horsetail (HT: −280′ to −20′), metaphase I (MTI: −10′ to 0′), meiosis I (MI: 10′ to 40′), metaphase II (MTII: 50′) and meiosis II (MII: 60′ to 70′). Column 3 showing closeup images of selected time frames from column 2. Numbers represent timing in minutes and scale bars equal 5 μm. (B) Durations of Rec8-GFP signal during metaphase I and meiosis I. (C) Quantitation of Rec8-GFP fluorescence intensity. More than 35 cells from at least two independent biological replicates movies were analyzed and representative image presented. Data presented in (B) and (C) represents mean timing and fluorescence intensities respectively. Error bars represent 95% class intervals.

During prefusion (PF: −310′ to −300′), we observed discrete and bright Rec8-GFP foci distributed across the nucleus (**Fig. 3A**). This pattern does not change during nuclear fusion (FS), horse-tailing (HT), and metaphase I (MTI) (FS to MTI: −290′ to 0′). From the beginning of horse-tailing until the end of metaphase I, Rec8 distribution, timing, and movement tracks those of H3 histone. A drastic change in signal intensity and nuclear pattern occurs as cells move from metaphase I (MTI) to meiosis I (MI). Within approximately 15 minutes (15.4′± 2.2′) of meiosis I, Rec8-GFP signal decreases until it becomes a single focus in each daughter nucleus (MTI to MI: 0′ to 20′), consistent with centromere localization (Watanabe and Nurse, 1999) (**Fig. 3B-C**). This focus remains until metaphase II (MI to MTII: 20′-50′) and disappears just before cells enter meiosis II (MII: 60′-70′). The complete loss of Rec8-GFP signal which precedes the final equational division of meiosis II, takes about 35 minutes (35.7′± 2.9′) from meiosis I (MI: 20′-50′) (**Fig. 3B**).

Our results agree with previous observations that Rec8 is detected in cells that have not yet undergone karyogamy (Doll et al., 2008; Watanabe and Nurse, 1999; Watanabe et al., 2001), which we observe are already in S phase. This is consistent with the nuclear organization function of Rec8 during recombination and metaphase I condensation (Ding et al., 2004; Ding et al., 2006; Ding et al., 2016). We have shown previously that disruption of DNA replication dynamics changes the relative timing of Rec8 in these events (Escorcia and Forsburg, 2017; Le et al., 2013; Mastro and Forsburg, 2014; Mastro et al., 2020), making this a particularly sensitive marker for meiotic progression.

Rec27, a meiotic recombination protein, is a part of the linear element (LinE) proteins that bind along the axes of homologous chromosomes to regulate chromosome pairing, programmed double strand breaks (DSB), and genetic recombination (Davis et al., 2008; Ellermeier and Smith, 2005; Phadnis et al., 2015; Sakuno and Watanabe, 2015). We used heterothallic strains containing homozygous Rec27-GFP and Hht1-mRFP and followed them from prefusion to meiosis II (Fig. 4). In early events, prior to nuclear fusion (PF), we observed a diffused, pan-nuclear Rec27-GFP signal which later turned into discrete puncta (PF: −210′ to −200′) (**Fig. 4A**). As cells entered karyogamy, the number and signal intensity of these Rec27-GFP foci increased (FS: −190′). During horse-tailing, the size of Rec27-GFP foci increased and changed into long linear structures (HT: −180′to −40′). As cells approach metaphase I, we observed gradual removal of those linear structures, converting into smaller foci with reduced signal intensity and number. This is followed by the complete elimination of Rec27-GFP signal (HT: −30′ to −0′). The disappearance of linear element signal happens about 20 minutes (19.1′± 3.2′) prior to the onset of metaphase I (**Fig. 4B**). We did not observe Rec27- GFP signal in metaphase I or following meiosis I and II (MTI to MII: −10′ to 60′) (**Fig. 4A, C**).

**Figure 4.**
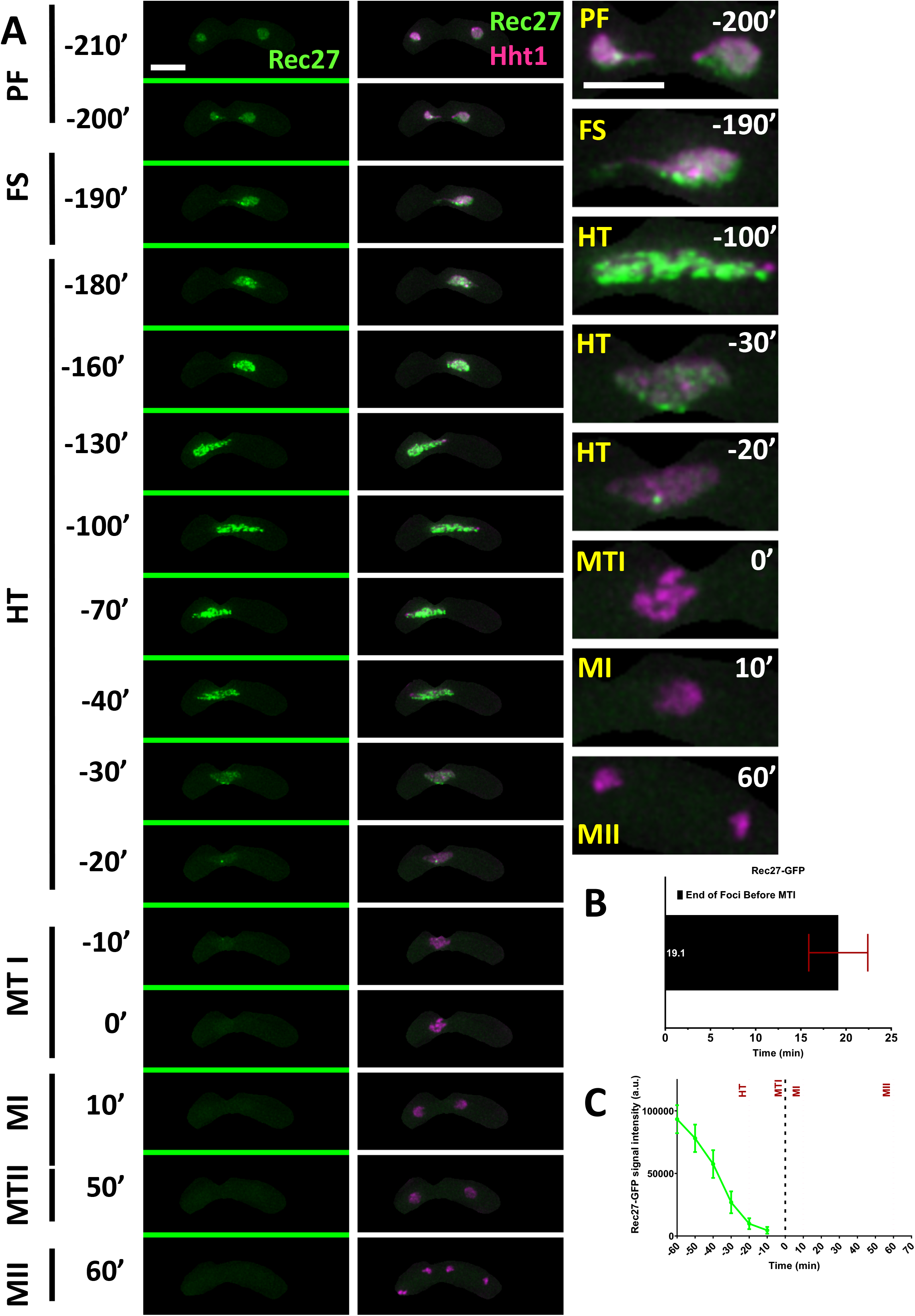
Nuclear dynamics of Linear element protein, Rec27. (A) Time-lapse images of meiotic cells carrying Rec27-GFP and Hht1-mRFP (7777 × 7779) were captured every 10 minutes for 8 hours and selected frames are shown. Panel (column 1: Rec27-GFP) and (column 2: merged Rec27-GFP Hht1-mRFP) show linear element dynamics during different meiotic events covering prefusion (PF), karyogamy (FS), horsetail (HT), metaphase I (MTI), meiosis I (MI), metaphase II (MTII) and meiosis II (MII). Numbers represent timing in minutes and scale bars equal 5 μm. (B) Durations of Rec27 foci elimination before metaphase I. (C) Quantitation of Rec27-GFP fluorescence intensity. More than 35 cells from at least two independent biological replicates movies were analyzed and representative image presented. Data in (B) and (C) represents mean timing and fluorescence intensities respectively. Error bars represent 95% class intervals.

Similar to our observation with Rec8-GFP and S-phase specific markers, the linear element protein Rec27-GFP is first observed in pre-fusion nuclei. Given our data with Pcn1 and Tos4, we infer that the Rec27 protein accumulates in the nucleus during pre-meiotic S phase and it may have some direct or indirect role in DNA replication. This observation is consistent with gene expression data suggesting Rec27 is expressed before premeiotic S phase (Mata et al., 2002). Previously, using nuclear spreads, it has been shown that Rec27 eventually forms distinct structures associated with linear elements (Davis et al., 2008). We observe that Rec27 foci begin to disappear as cells transition from late horse-tailing into metaphase I. This observation reflects the end of Rec27 function in chromosome alignment and recombination (Bähler et al., 1993; Davis et al., 2008; Ellermeier and Smith, 2005; Phadnis et al., 2015; Sakuno and Watanabe, 2015). Since disruption of Rec27 dynamics is associated with chromosome mis-segregation (Escorcia and Forsburg, 2017), the time-frame before nuclear condensation suggests a promising stage to study LinE proteins in the process of crossing-over and chiasma resolution (Ding et al., 2004; Sharif et al., 2002).

The meiotic inner centromere protein, Shugoshin (Sgo1) along with phosphatase protein 2A (PP2A) helps to protect Rec8 from proteolytic degradation at the centromere prior to meiosis I (Ishiguro et al., 2010; Kitajima et al., 2004; Riedel et al., 2006). We observed meiosis in cells carrying homozygous alleles of Sgo1-GFP and Hht1-mRFP. Sgo1-GFP signal emerges when nuclear oscillation slows down in late horse-tailing. At this stage, Sgo1-GFP is diffused throughout the nucleus (pan-nuclear) but then resolves into discrete foci. These foci and nuclear signal remain visible for another 50-55 minutes before disappearing during the transition between metaphase I (MTI) to meiosis I (HT to MTI: −60′ to 0′) (**Fig. 5A**). These punctate foci disappear within 11 minutes (11.4′± 1.1′) of the metaphase I to meiosis I transition (MTI to MI: 0′ to 10′) (**Fig. 5B**). We did not observe any Sgo1 foci or fluorescence signal after cells transitioned into meiosis I and could only capture the background (MTI to MII: 0′ to 70′) (**Fig. 5A, C**). The emergence of Sgo1-GFP foci, approximately an hour before metaphase I are consistent with its role as protector of centromeric Rec8 degradation prior to meiosis I (Ishiguro et al., 2010; Kitajima et al., 2004; Kitajima et al., 2006). Our data agree with other studies showing Rec8-GFP is removed from chromosome arms but not the centromere, where Sgo1 localization prevents cleavage (Ishiguro et al., 2010; Mastro et al., 2020).

**Figure 5.**
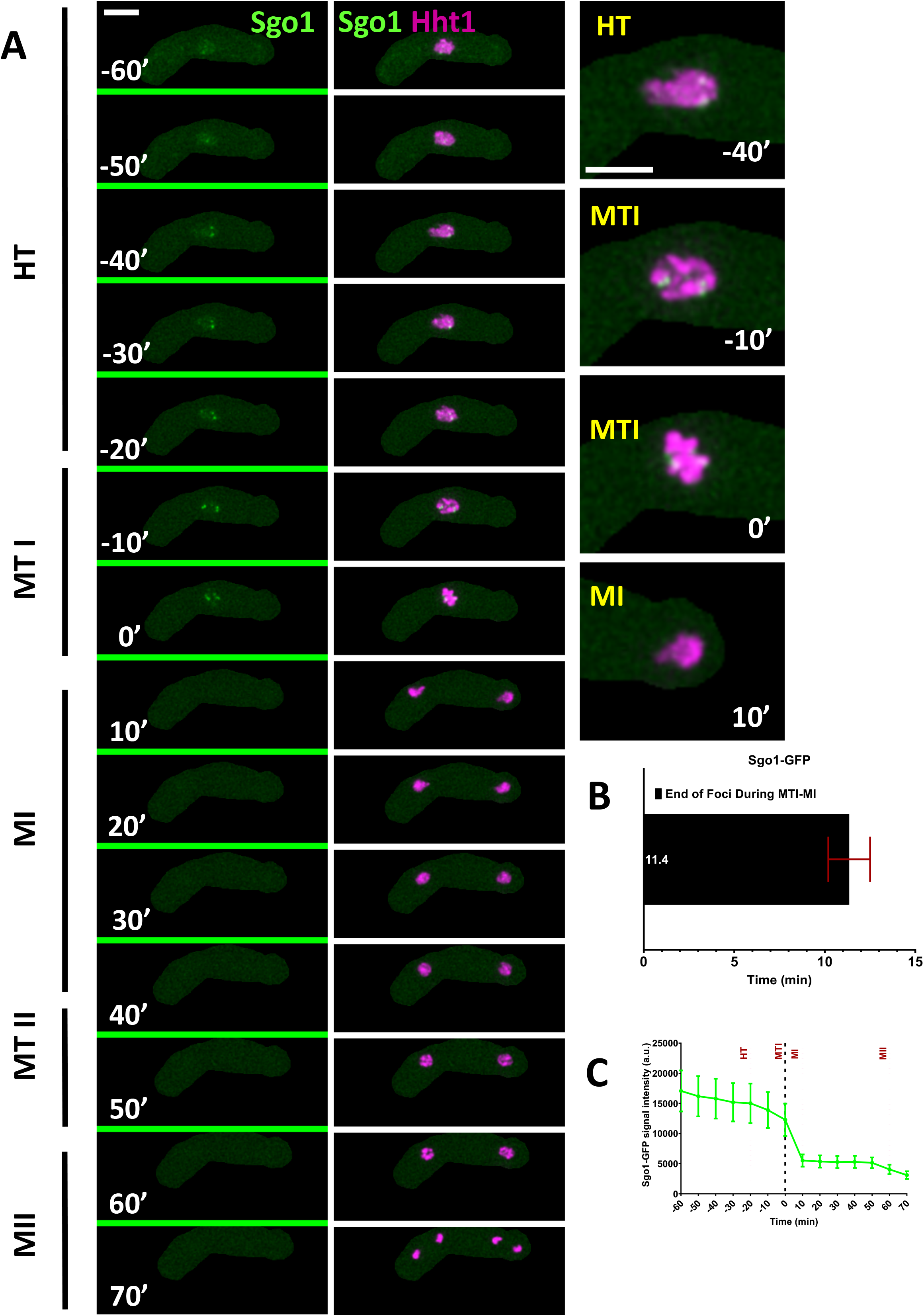
Nuclear dynamics of Shugoshin (Sgo1) protein. (A) Time-lapse images of meiotic cells carrying Sgo1-GFP and Hht1-mRFP (7864 × 7865) were captured every 10 minutes for 8 hours and selected frames are shown. Panel show nuclear dynamics during different meiotic events covering horsetail (HT) to meiosis II (MII). Numbers represent timing in minutes and scale bars equal 5 μm. (B) Durations of Sgo1-GFP foci disappearance during metaphase I. (C) Quantitation of Rec27-GFP fluorescence intensity. More than 35 cells from at least two independent biological replicates movies were analyzed and representative image presented. Data in (B) and (C) represents mean timing and fluorescence intensities respectively. Error bars represent 95% class intervals.

Moa1 (monopolin), a meiosis specific kinetochore protein, is required to establish monopolar attachment of sister chromatids during meiosis I (Yokobayashi and Watanabe, 2005). It associates with Rec8 at the centromere and helps to stabilize Sgo1, thereby ensuring protection of centromeric cohesion, which is crucial for separation of homologous chromosomes (Galander et al., 2019; Hirose et al., 2011; Sakuno et al., 2009; Yokobayashi and Watanabe, 2005). Similar to Sgo1-GFP, we followed zygotic meiosis in cells carrying homozygous alleles of Moa1-GFP and Hht1-mRFP (**Fig. 6**). We did not see any GFP signal in early stages of PF, FS or early HT (data not shown). However, we saw discrete Moa1-GFP foci appear during later horse-tailing (HT: −60′) **(Fig. 6A)**. Moa1-GFP foci remain visible for about 48 minutes (48.6 ± 3′) during late horse tailing and early metaphase I (HT to MTI: −60′ to 0′) (**Fig. 6A-C**). The foci are initially dispersed around the nucleus and then cluster together at discrete spots at the poles of the condensing nuclei (MTI: −10′ to 0′). During metaphase I (MTI: 0′). Moa1-GFP forms a bright structure at the poles of the separating nuclei, presumably at the kinetochore, with a less intense dot just inside the nuclear mass. Within 14 minutes (14′± 2′) of the metaphase I to meiosis I transition (MTI to MI: 0′-10′), we observe a complete loss of Moa1-GFP foci. We did not see any signal post anaphase I (MI to MII: 10′ to 70′). Moa1 is reported to localize in the central core of the centromere, where it interacts with Rec8, ensuring that sister chromatids face the same orientation prior to homolog separation (Kim et al., 2015; Miyazaki et al., 2017; Yokobayashi and Watanabe, 2005). The role of Moa1 in mono-orientation is evident from the separation of foci clusters that occurs as cells approach metaphase I. This signal pattern is similar to that of Sgo1-GFP, thereby lending additional support to their functional association (Kitajima et al., 2004; Kitajima et al., 2006).

**Figure 6.**
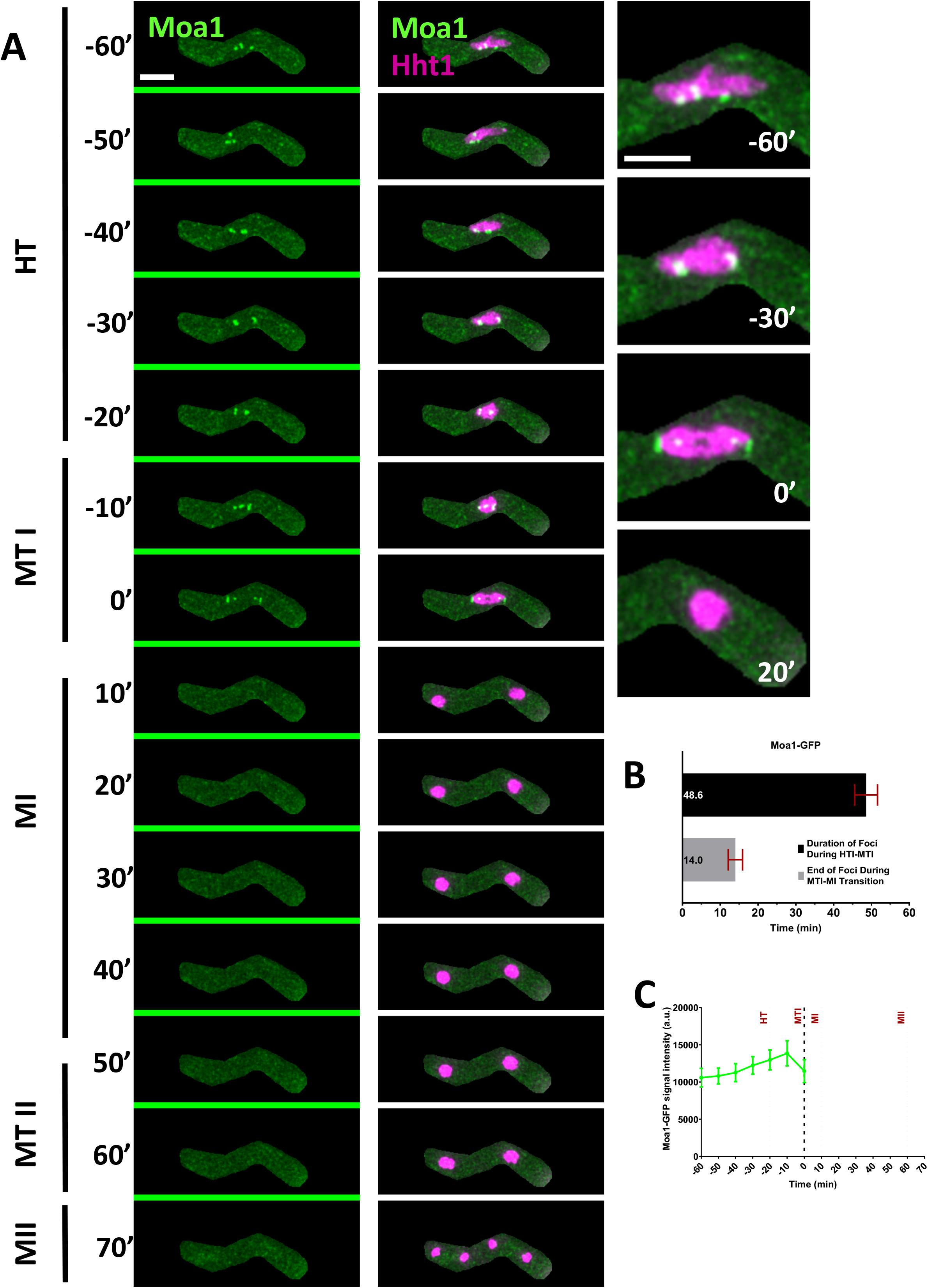
Nuclear dynamics of a kinetochore regulator Moa1. (A) Time-lapse images of meiotic cells carrying Moa1-GFP and Hht1-mRFP (8047 × 8048) were captured every 10 minutes for 8 hours and selected frames are shown. Panel show nuclear dynamics during different meiotic events covering horsetail (HT) to meiosis II (MII). Numbers represent timing in minutes and scale bars equal 5 μm. (B) Durations of Moa1-GFP foci during horsetails and metaphase I. (C) Quantitation of Moa1-GFP fluorescence intensity. More than 35 cells from at least two independent biological replicates movies were analyzed and representative image presented. Data in (B) and (C) represents mean timing and fluorescence intensities respectively. Error bars represent 95% class intervals.

### Recombination and Repair

Rad11 (Ssb1) is the fission yeast orthologue of replication protein A (RPA), which binds to single stranded DNA (ssDNA) resulting from replisome progression in DNA synthesis and processing of DSB during DNA repair and homologous recombination (Escorcia and Forsburg, 2017; Mastro and Forsburg, 2014; Parker et al., 1997; Sabatinos et al., 2012). To observe signal dynamics of recombination and repair proteins, prior to and during the meiotic nuclear divisions, we examined cells with fluorescently marked RPA (Rad11-CFP), Rad52 (Rad52-YFP) and Hht1(Hht1-mRFP). During prefusion (PF: −270′), we observed a pan-nuclear Rad11-CFP signal, which later forms discrete foci distributed across the nucleus from karyogamy (FS) until late horse-tailing (HT) stages, when the nuclear oscillation starts slowing down (PF to HT: −270′ to −50′ (**Fig. S2A**)). As cells progressed towards the end of the horse tailing, the number of foci and overall fluorescent intensity decreased and is finally lost before metaphase I (HT to MTI: −70′ to 0′), (**Fig. 7A, E**). The average mean time of the disappearance of these discrete foci, prior to the metaphase I, is about 20 minutes (21′± 2.3′) (**Fig. 7C)**. From metaphase I onwards, we did not observe any foci. After meiosis I, and for the next 53 minutes (53.4′± 1.8′), Rad11-CFP signal dissipates in the newly formed nuclei and finally ends before meiosis II (MI to MII: 0′ to 70′) (**Fig. 7C, E).**

**Figure 7.**
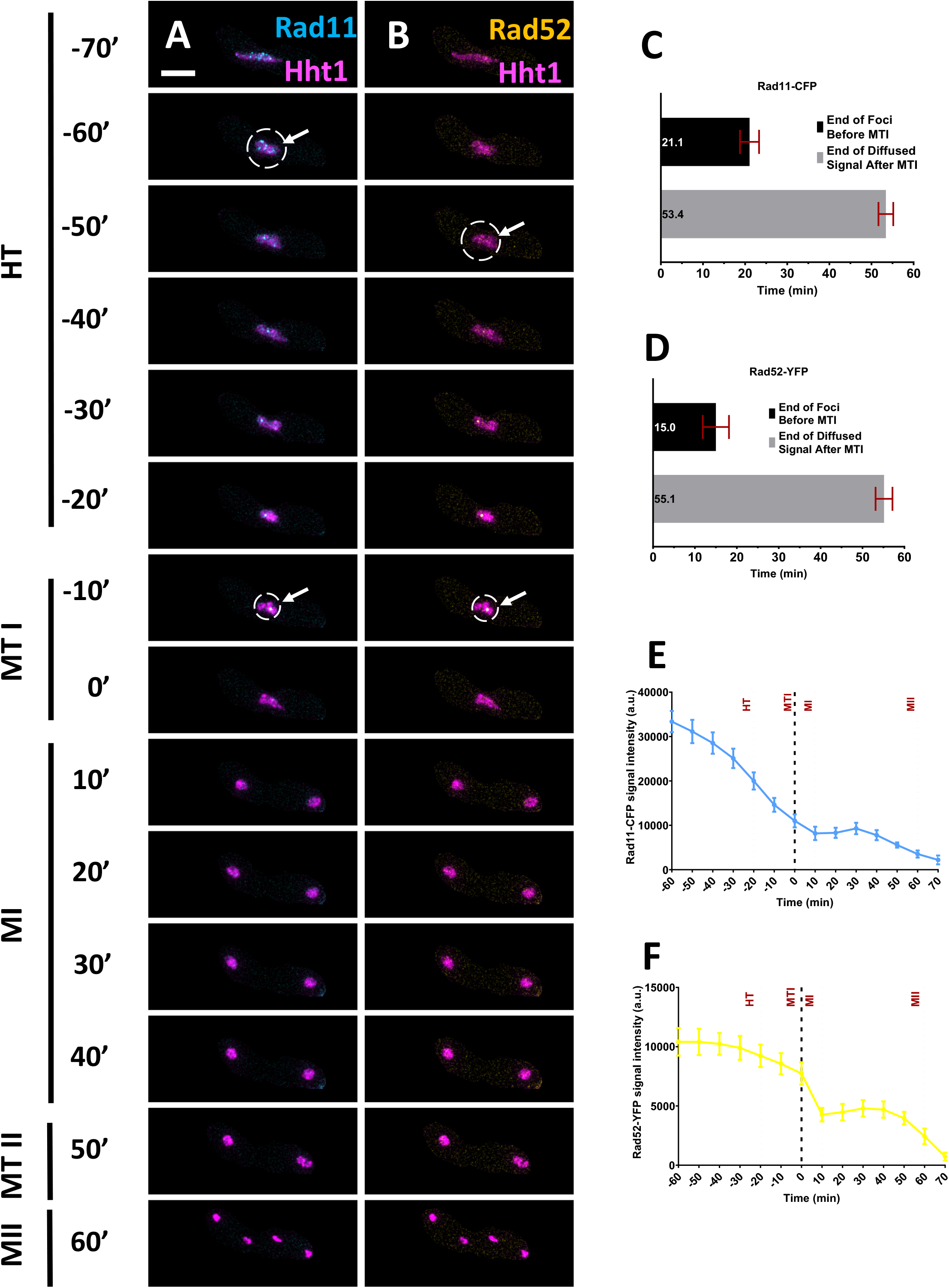
Nuclear dynamics of Rad11 (RPA) and Rad52 proteins. (A) Time-lapse images of meiotic cells carrying Rad11-CFP, Rad52-YFP and Hht1-mRFP (7327 × 7328) were captured every 10 minutes for 8 hours and selected frames are shown. Panels (A) merged Rad11-CFP, Hht1-mRFP and (B) merged Rad52-YFP, Hht1-mRFP, are showing nuclear dynamics during different meiotic events encompassing horsetail (HT) to meiosis II (MII). Numbers represent timing in minutes and scale bars equal 5 μm. White arrow and circle showing presence of Rad11 and Rad52 foci signals. More than 35 cells from at least two independent biological replicates movies were analyzed and representative image presented. (C-D) Showing durations of Rad11(C) and Rad52 (D) foci signal loss, prior to metaphase I stage. (E, F) Quantitation of Rad11-CFP (E) and Rad52-YFP (F) fluorescence intensity. More than 35 cells from at least two independent biological replicates movies were analyzed and representative image presented. Data in (C-D and E-F) represents mean values of timing and fluorescence intensities respectively. Error bars represent 95% class intervals.

Rad52, a DNA recombination protein, acts with RPA and Rad51 to regulate recombination, which facilitates reductional division of the homologous chromosomes in meiosis I (Murayama et al., 2013; Muris et al., 1997; Octobre et al., 2008; van den Bosch et al., 2001). In our analysis, we did not see any Rad52-YFP signal during the early stages (PF to HT: −270′ to −50′) including prefusion, karyogamy, or early horse-tailing (**Fig. S2B)**. We noticed the emergence of Rad52 signal during late horse-tailing, the time when cells are undergoing recombination and repair events. We observed discrete punctate foci during late horse-tailing. These decrease in intensity and diffuse as the cells enter metaphase I (HT to MTI: −50′ to 0′) (**Fig. 7B**). Rad52 foci lasts about 15 minutes (15′± 3′) prior to metaphase I, which is 5 minutes later than Rad11 (**Fig. 7D**). We also noticed that, prior to and during metaphase I, some of the Rad11-CFP foci colocalizes with Rad52-YFP foci (HT to MTI: −50′ to −10′) (**Fig. 7A, B**). Both Rad52 and RPA puncta are resolved by the time cells proceed through the meiosis I division. We observed a similar, diffuse nuclear Rad52 signal for about 55 minutes (55′± 1′) after meiosis I until it finally disappears **(Fig. 7D, F)**.

These results indicate that proteins involved in replication, and repair span the timing from pre-fusion to metaphase I. Rad11, a large subunit of RPA, is associated with DNA synthesis, repair, and recombination (Parker et al., 1997). The presence of discrete Rad11-CFP foci across the nucleus during prefusion resemble the nuclear dynamics of Pcn1, Tos4 and Rec8. This result is consistent with premeiotic S-phase initiating during the prefusion stage. The nuclear dynamics of Rad11 during late horse-tailing, where CFP foci starts to decrease in number and intensity resembles those of Rec27, suggesting the timing of recombination resolution.

Rad52 is a recombination mediator that either promotes Rad51 or antagonizes Dmc1 binding to RPA-coated presynaptic filaments during recombination (Murayama et al., 2013; Muris et al., 1997; Octobre et al., 2008). Which recombinase follows Rad52 association with RPA largely determines recombination dynamics in meiotic prophase (Murayama et al., 2013). Our observations show that Rad52-YFP signal is diffused until late horse-tailing, when it forms punctate foci. Timing of Rad52 foci emergence and its colocalization with Rad11 foci during late horse-tailing suggest the timing of recombination and repair activities, which finally concludes before metaphase I, when both signals disappear.

### Spindle and spindle checkpoint dynamics

The final group of proteins we studied are involved in microtubule dynamics and the spindle checkpoint. The spindle protein Atb2, a component of α-tubulin, is a constituent of cytoplasmic microtubule organization and is actively involved in establishment and maintenance of cell polarity and cell shape (Adachi et al., 1986; Radcliffe et al., 1998). We crossed strains heterozygous for Atb2-mCherry) (Yamagishi et al., 2008) and Hht1-GFP, and imaged meiotic progression from pre-fusion to meiosis II. Initially, during prefusion, we observed a bundle of microtubules (Atb2-mCherry) on one end of the cell and the Hht1-GFP signal on the other (PF: −220′ to −200′) (**Fig.S3A**). As cells move to karyogamy, the bundle of microtubules approach the Hht1-GFP nucleus (FS: −200′ to −180′). Nuclear oscillations initiate with horse-tailing (HT: −160′). The microtubules show repeated expansion and contraction as they move across the cell (HT: −160′ to −40′; **Fig. S3**) and (HT: −60′ to −40′; **Fig. 8A**). At the end of horse-tailing microtubules briefly disappear, and the Atb2-mCherry signal returns as a spindle prior to metaphase I (HT: −50′ to −20′) (**Fig. 8A, B)**; this gradually increases in length and fluorescence signal intensity during the prophase I stage of meiosis I (**Fig. 8A-B, Fig. S3B**). At metaphase I, which lasts for about 12 minutes (12.3′± 1.5′), there is an increase in Atb2-mCherry signal intensity with slight change in spindle lengths (MTI: 0′) **(Fig. 8A, Fig. S3B**). At anaphase I (MI), microtubules elongate outwards and reached to the cell’s tip. The elongated microtubules carry newly separated daughter nuclei to the opposite ends of the cells and the process lasts for about 11 minutes (10.9′± 1′) (MI: 10′ to 20′) (**Fig. 8A, D, Fig. S3B)**. Next, for about 23 minutes, microtubules undergo depolymerization and reappearance of Atb2-mCherry signals showing completion of meiosis I and the beginning of prophase II of meiosis II stage (MI: 40′) (**Fig. 8A, B, Fig. S3B)**. Finally, at the end of meiosis II the Atb2-mCherry signals disappeared and four completely separated daughter nuclei formed (MII: 70′). The duration of metaphase II was approximately 12 minutes (12.3′± 1.5′) while anaphase II lasted for about 14 minutes (14.6′± 1.8′) (**Fig. S3B)**. The spindle length and signal intensity in meiosis I and II are proportional to the respective nuclear mass (hht1-GFP) (**Fig. 8B)**.

**Figure 8.**
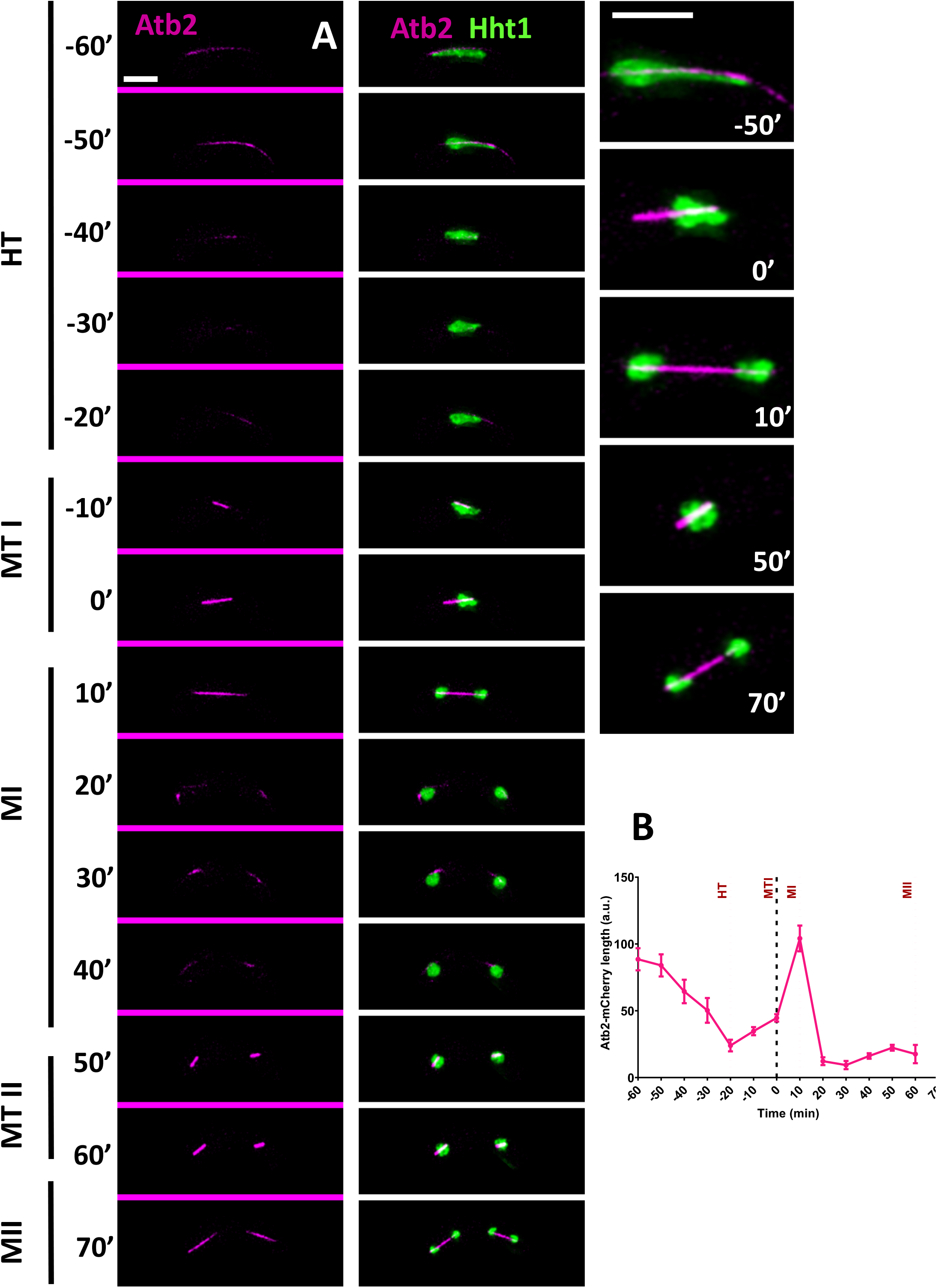
Meiotic spindle (Atb2-mCherry) dynamics. (A) Time-lapse images of unperturbed zygotic meiotic cells carrying fluorescently tagged heterozygous histone (Hht1-GFP) and tubulin (Atb2-mCherry) (8055 × 8203) were captured every 10 minutes for 8 hours and selected frames are shown. Panel show spindle dynamics covering different meiotic events from horse-tailing (HT) to meiosis II (MII). Column 3 are a closeup images of selected time frames from middle panel (merged Atb2-mCherry and Hht1-GFP). Numbers represent timing in minutes and scale bars equal 5 μm. More than 35 cells from at least two independent biological replicates movies were analyzed and representative image presented. (B) Quantitation of the length of microtubule. Data represents average length of microtubules. Error bars represent 95% class intervals.

In *S. pombe* meiosis, microtubules play crucial roles from early stages including fusion and horse tailing (Katsumata et al., 2016). Our prefusion and karyogamy stage data are consistent with previous work showing a bundle of microtubules mobilizing and forming a X-shaped structure (Ding et al., 1998). Data from meiosis I and II stages are in alignment with previous reports showing three distinct steps in spindle elongation: prophase, metaphase characterized by constant spindle length with increased signal intensity, and anaphase, when microtubule elongates further leading to the edge of the cell with slight change in the signal intensities (Miyazaki et al., 2017; Nabeshima et al., 1998; Sakuno et al., 2011; Yamamoto et al., 2008).

Aurora-B kinase (*Sp*Ark1) is a serine/ threonine kinase which ensures phosphorylation of substrates involved during chromosome condensation, activation of spindle assembly checkpoints and correction of erroneous kinetochore-microtubule attachments (Hauf et al., 2007; Kawashima et al., 2007; Matsuhara and Yamamoto, 2016; Petersen et al., 2001; Sakuno et al., 2011). To examine its dynamics in zygotic meiosis with reference to microtubules, we used heterothallic strains containing Ark1-GFP and Atb2-mCherry. We did not see any Ark1-GFP signal during early events including prefusion, karyogamy or early horse tailing stages (data not shown). The first Ark1-GFP signal was observed during the later stage of horse-tailing (HT: −20′), just prior to the beginning of metaphase I (**Fig. 9A**). Initially, for about 9 minutes (8.9′± 1.1′), it appeared as a discrete bright punctate focus, located in the central region of Atb2 (**Fig. 9A, Fig. S4A)**. At metaphase I, the number and fluorescence intensity of Ark1-GFP foci increased drastically and were distributed across the length of the spindle except on the two poles (MTI: −10′ to 0′) (**Fig. 9A, Fig. S4B)**. This scattered pattern of Ark1-GFP foci signal lasts for about 13 minutes (13.7′± 1.6′) (**Fig. S4A)**. At anaphase I (MI: 10′), Ark1-GFP signal moves towards the midzone and localizes to the midbody of the extended spindles for about 14 minutes (14.3′± 2′). At this stage, the fluorescence signal intensity of Ark1-GFP reached a maximum **(Fig. S4B)**. After completing anaphase I, Ark1-GFP briefly disappears for about 13 minutes (13.4′± 1.7′) before showing similar dynamics in meiosis II: reappearance of Ark1-GFP signal during prophase II for 8 minutes (8.3′± 1.3′), redistribution to the spindles length during metaphase II for 17 minutes (17.7′± 1.5′) and back to the midbody on the spindles during anaphase II for 9 minutes (8.9′±1.1′) following the similar pattern as observed in meiosis I (MI-MII: 30′ to 60′) (**Fig. 9A, Fig. S4A, B)**. Ark1-GFP signal disappears after the completion of sister chromatid segregation at meiosis II.

**Figure 9.**
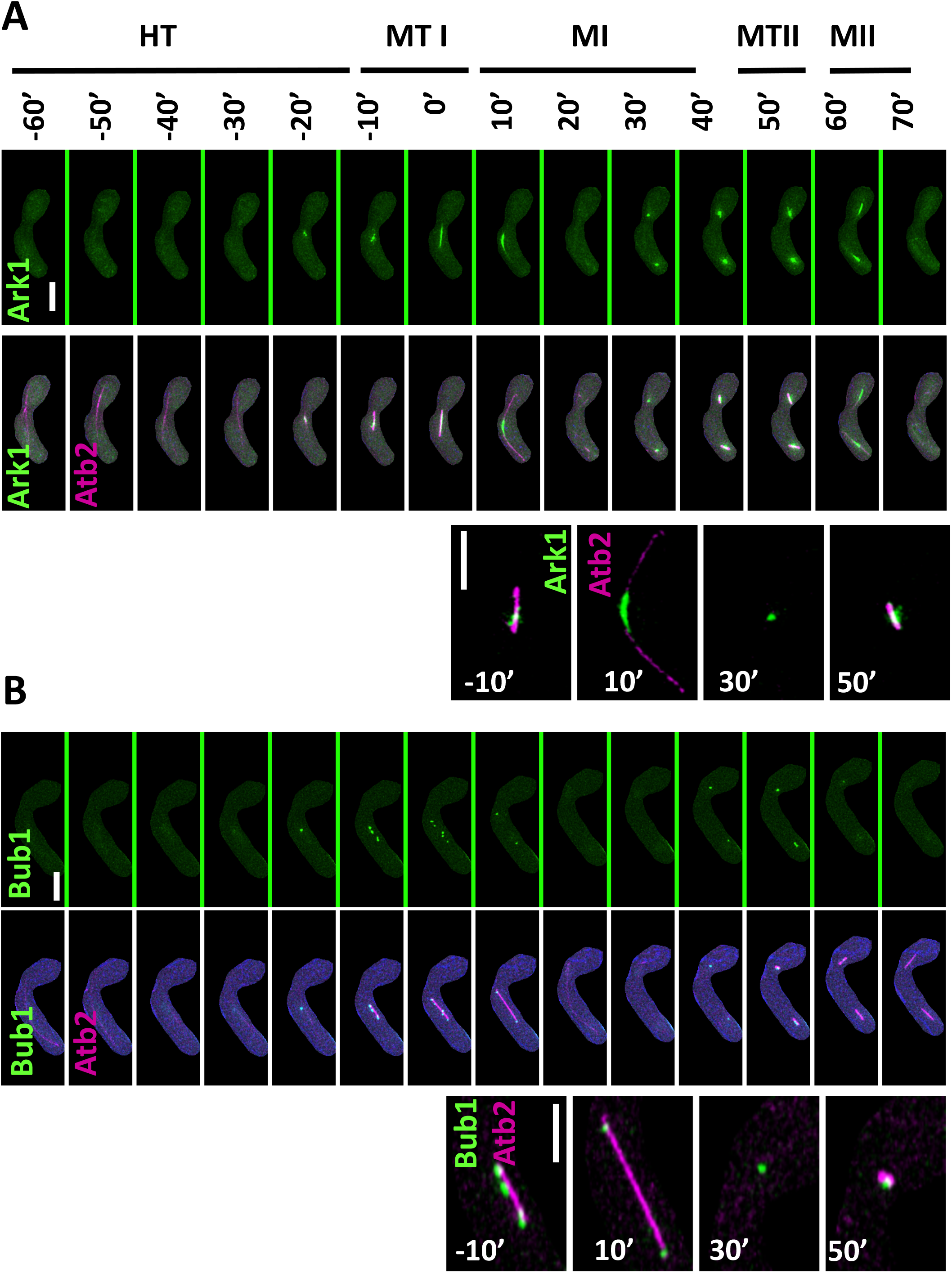
Nuclear dynamics of Aurora B (Ark1) and Bub1 kinase. Live-cell images of unperturbed zygotic meiotic cells carrying fluorescently tagged (A) Aurora B kinase (Ark1-GFP)& microtubule (Atb2-mCherry) (8053 × 8056) and (B) Bub1 kinase (Bub1-GFP)& microtubule (Atb2-mCherry) (8055 × 8067). Panels showing nuclear dynamics during different meiotic events encompassing horse-tailing (HT: −60′) up to meiosis II (MII: −70′). Row 3^rd^ and 6^th^ are the images of selected time frames from row 2^nd^ and 5^th^ respectively. Numbers represent timing in minutes and scale bars equal 5 μm. More than 35 cells from at least two independent biological replicate movies and representative image presented. Quantitative data presented in figure S4.

Our results showing appearance of discrete bright Ark1-GFP foci, near mid spindle region, agree with previous reports (Hauf et al., 2007; Petersen et al., 2001; Sakuno et al., 2011) and are consistent with enrichment on centromeres which might be aligned on the spindle. Sgo1 shows similar dynamics during prometaphase and metaphase and Ark1 colocalizes with Sgo1 at centromeres and SPBs (Kawashima et al., 2007; Petersen et al., 2001). The localization of Aurora kinase on peri-centromeres during prometaphase has been confirmed using ChIP (Morishita et al., 2001). The durations and dynamics of Ark1-GFP signals are in alignment with spindle polymerization suggesting its crucial role in proper spindle maintenance. Our Ark1 dynamics of meiosis II aligned with meiosis I and mitotic cell cycles (Hauf et al., 2007; Petersen et al., 2001; Sakuno et al., 2011; Salas-Pino and Daga, 2019).

Next we examined an essential component of spindle assembly checkpoint, Bub1, a serine/threonine kinase, required for the protection of meiotic centromeric cohesion (Bernard et al., 2001; Marston and Wassmann, 2017). Previously, its dynamics were examined in *pat1* synchronized meiosis but how it behaves in an unperturbed meiosis was not tested (Bernard et al., 2001; Miyazaki et al., 2017). Here, we used heterozygous cells carrying GFP tagged Bub1 (Bub1-GFP) and mCherry tagged Atb2 (Atb2-mCherry) and followed the zygotic meiosis (Fig. 9B). We did not observe any Bub1-GFP signal until late horse-tailing stage where we see a faint pan-nuclear signal (HT: −40′) which later form a bright, discrete focus (HT: −20′). The pattern remains visible for about 18 minutes (18.3′± 2.3′) during late horse-tailing stage (Fig S4C). As cells reached metaphase I, the single Bub1-GFP focus split into a maximum of six foci distributed across the metaphase I spindle (**Fig. 9B**). Bub1-GFP signal intensity increases drastically and reached to the maximum which lasts for about 11 minutes (11.4′± 1.2′) (MTI: 0′) (**Fig. 9B, Fig. S4C, D)**. At meiosis I, the signal across the spindle diffused and coalesced at a single focus on each daughter nucleus. Previous work suggests that Bub1-GFP enrichment at the ends of the spindle represents its localization at the kinetochores (MI: 10′) (Miyazaki et al., 2017). This stage lasts for 10 minutes (10.3′± 0.5′) (**Fig. S4C**). As cells complete anaphase I, the Bub1-GFP signal disappears and returns after 20 minutes (20.9′± 1.7′) post anaphase I, when again it appears in puncta for about 10 minutes (10.3′± 0.6′) (**Fig. 9B, Fig. S4C**). As cells enter metaphase II, the signal diffuses and distributes across the spindle in the same fashion as in metaphase I (MI-MII: 40′ to 60′). In this state, Bub1-GFP signals remain visible for 18 minutes (18.9′± 1.4′) before disappearing during anaphase II (**Fig. S8C**). We did not see any Bub1-GFP signal during the meiosis II. Bub1 plays an important role in centromeric cohesion protection and proper Sgo1 localization to the pericentromeric regions (Bernard et al., 2001; Kawashima et al., 2010; Kitajima et al., 2004). Bub1 also plays a crucial role in Ark1 localization to the centromere regions (Hauf et al., 2007; Kawashima et al., 2007; Kitajima et al., 2004). Our results show that Bub1 and Ark1, during prophase and metaphase stages, show similar timing and behavior. There is a difference during anaphase, when Ark1 leaves centromeric region and localizes to the midzone of spindle, while Bub1 remain attached to the centromeres. Our result also showed different Bub1 dynamics during meiosis I and II, as it remains attached to the centromere in meiosis I, but we could not see any signal during meiosis II. This supports previous findings of Bub1′s role in monopolar attachment (Hauf et al., 2007; Miyazaki et al., 2017).

### Concluding remarks

Fission yeast is an excellent model for meiotic progression. Typically, investigators describe the behavior of one or a few novel proteins during meiosis, but protocols vary from lab to lab. In many cases, investigators make use of the *pat1-114* temperature sensitive mutant (Beach et al., 1985; Iino and Yamamoto, 1985a; Iino and Yamamoto, 1985b; Nurse, 1985) which can induce a synchronous meiosis from haploid or diploid cells in response to temperature shift. However, *pat1* driven meiosis has notable differences from a normal zygotic meiosis, and these are exacerbated in haploid meiosis compared to diploids (e.g., (Bähler et al., 1991; Pankratz and Forsburg, 2005; Yamamoto and Hiraoka, 2003). Additionally, many studies employ “snapshots” of different timepoints, rather than continuous observations with live cell imaging that reveal dynamic and brief events. We have established a standardized protocol for imaging meiosis over time in normal homothallic or heterothallic strains, which we employed previously (Escorcia et al., 2019). In this study, we chose representative proteins involved in different stages of meiosis, and compared their signal intensity, localization, and dynamics against a standard panel of meiotic nuclear events including karyogamy, horsetail formation, meiosis I and meiosis II divisions. Based on these analyses, we present a reference that defines dynamic behavior of proteins during meiotic DNA synthesis, genomic fusion, chromosome alignment, genetic recombination, metaphase, and meiosis, using consistent conditions (**Fig. 10**). This live-cell microscopy approach provides information about the change of protein abundance and localization during meiotic differentiation and establishes a common template to facilitate comparisons of different proteins. This should prove particularly helpful in mutant backgrounds to define the extent of perturbations. By capturing the timing of different meiotic events in the context of nuclear change, we provide a general atlas of reliable markers that can be used to study dynamics and disruptions of meiotic processes in *S. pombe.*

**Figure 10.**
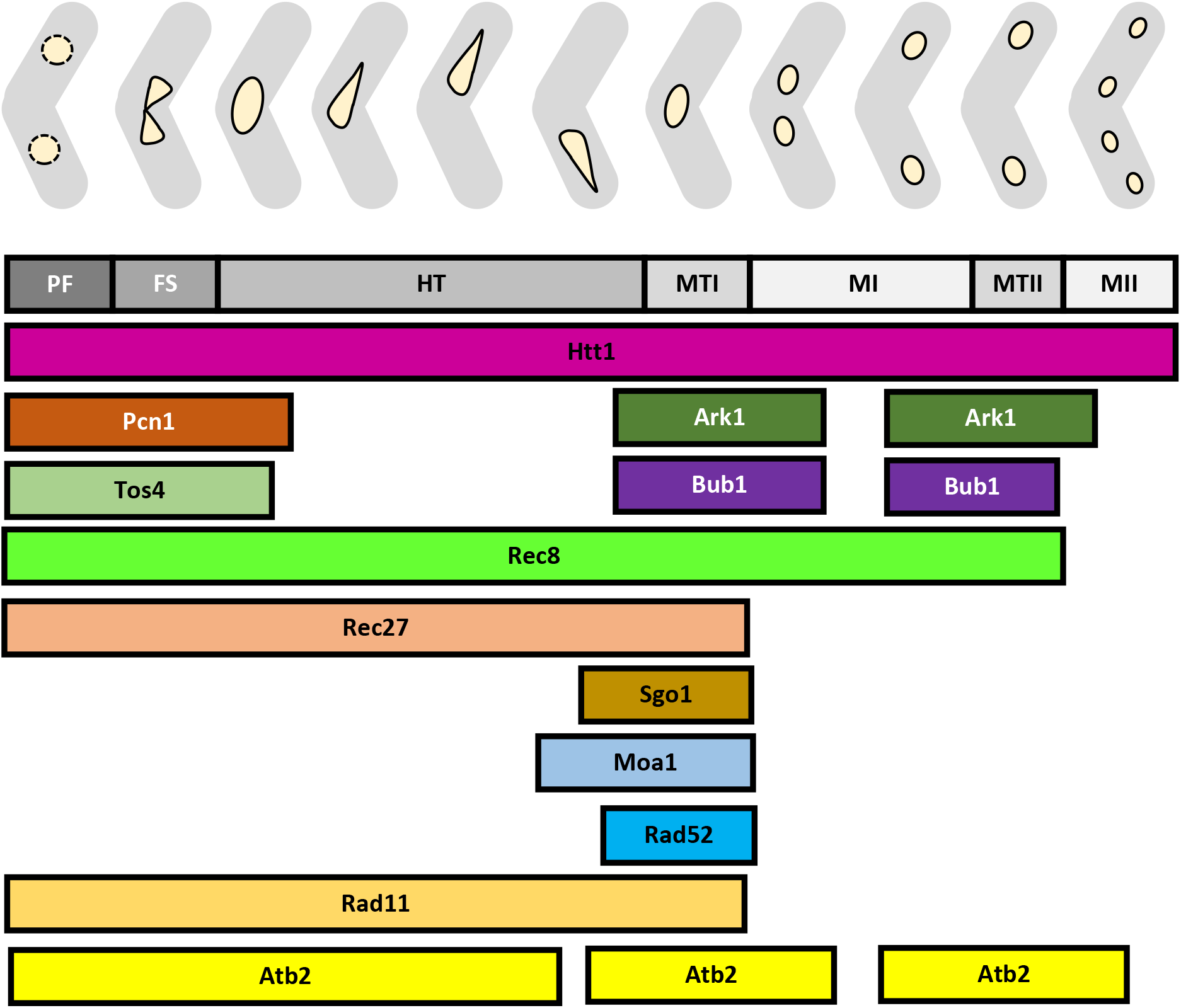
Meiotic signposts. Model presenting a visual atlas of reliable meiotic proteins that can be used to study the nuclear dynamics and disruptions of different meiotic processes in *S. pombe.* Meiotic signposts encompass events including prefusion (PF), Karyogamy (FS), horse-tailing (HT), metaphase I (MTI), meiosis I (MI), metaphase II (MTII) and meiosis II (MII). Respective timing of presence of different meiotic proteins during described meiotic signposts are presented.

## Materials and Methods

### Yeast strains, culture and cell growth

General fission yeast culture condition, media compositions and strain constructions are used as described in (Sabatinos and Forsburg, 2010). Different fluorescently tagged *S. pombe* strains used in this study are listed in the supplementary Table S1. For live cell imaging, heterothallic *(h+* and *h-)* cells were initially grown in 5 ml YES media at 32°C for 12-16 hours. 100- 250 μl starter culture transferred to 5 ml EMM media containing appropriate supplements and grown at 32°C until the culture reached a late log stage (OD_595_ =~0.8). Late log phase cells harvested at 3,000 rpm for 5 minutes. To remove any remaining nutrient from the culture, pellets were washed twice with Malt Extract (ME) media. Finally, heterothallic pellets resuspended in 10 ml ME media and incubated at 25°C for 12 to 16 hours at 50-60 rpm. From this starved mating culture, 1 ml culture harvested in microfuge tubes and used for live cell microscopy.

### Fluorescence live-cell microscopy

Live cell imaging of meiotic events was performed as described in (Escorcia and Forsburg, 2017; Escorcia et al., 2019). Briefly, 1 ml mating culture harvested at 5,000 rpm for 30 seconds. Pellets were resuspended in 250 μl ME media. 10 μl of cell suspension spread on top of 2% agarose pad made with liquid sporulation media and sealed with VaLaP (Vaseline/ Lanolin/ Paraffin in a ratio 1:1:1 by weight). Live cell imaging was performed at 25°C (pre-calibrated chamber) on a Delta Vision Microscope (Applied Precision, GE Healthcare, Issaquah, WA) equipped with Olympus 60×/1.40 Plan-Apo objective lens, solid-state illuminator, and 12-bit Photo metrics CoolSNAP_HQ2 Charged-coupled device (CCD) camera. Different filter sets and exposure timings were used to excite and detect fluorescent proteins (Table 2). 13 optical *z*-sections of 0.5 μm step size were acquired for each field at 10-minutes intervals over 8 hours. Images were acquired, deconvolved and all z-stacks projected into a single-plane as maximum intensity projection by SoftWoRx Version 5.5.1 (GE, Issaquah, WA) software. Finally, the projected fluorescence images were fused with transmitted light images. Downstream processing and image analysis were done using Fiji (ImageJ) an open source image analysis software (Schindelin et al., 2012). Detailed description of image analysis is described in (Escorcia and Forsburg, 2017; Escorcia et al., 2019).

### Statistical Analysis

For different marker metrics across time and timing of individual events, the mean with 95% CI is presented. For comparisons among time points per marker, significance was evaluated using an ordinary One-Way ANOVA (for matched data) followed by Tukey’s post hoc test. GraphPad Prism 8.4.2 was used for statistical analysis and graphical representations of the data.

## Acknowledgements

We thank Julie Cooper, Gerry Smith, and Yoshi Watanabe for sharing strains. The members of the lab for helpful comments. Marc Green and Tara Mastro for guidance and mentorship. Rosalind Elsie Franklin and Percy Lavon Julian for scientific inspiration. Funding from NIGMS R35 GM118109 (SLF) and T32AG052374 (WE).

## Supplementary Figures

**Figure S1.**
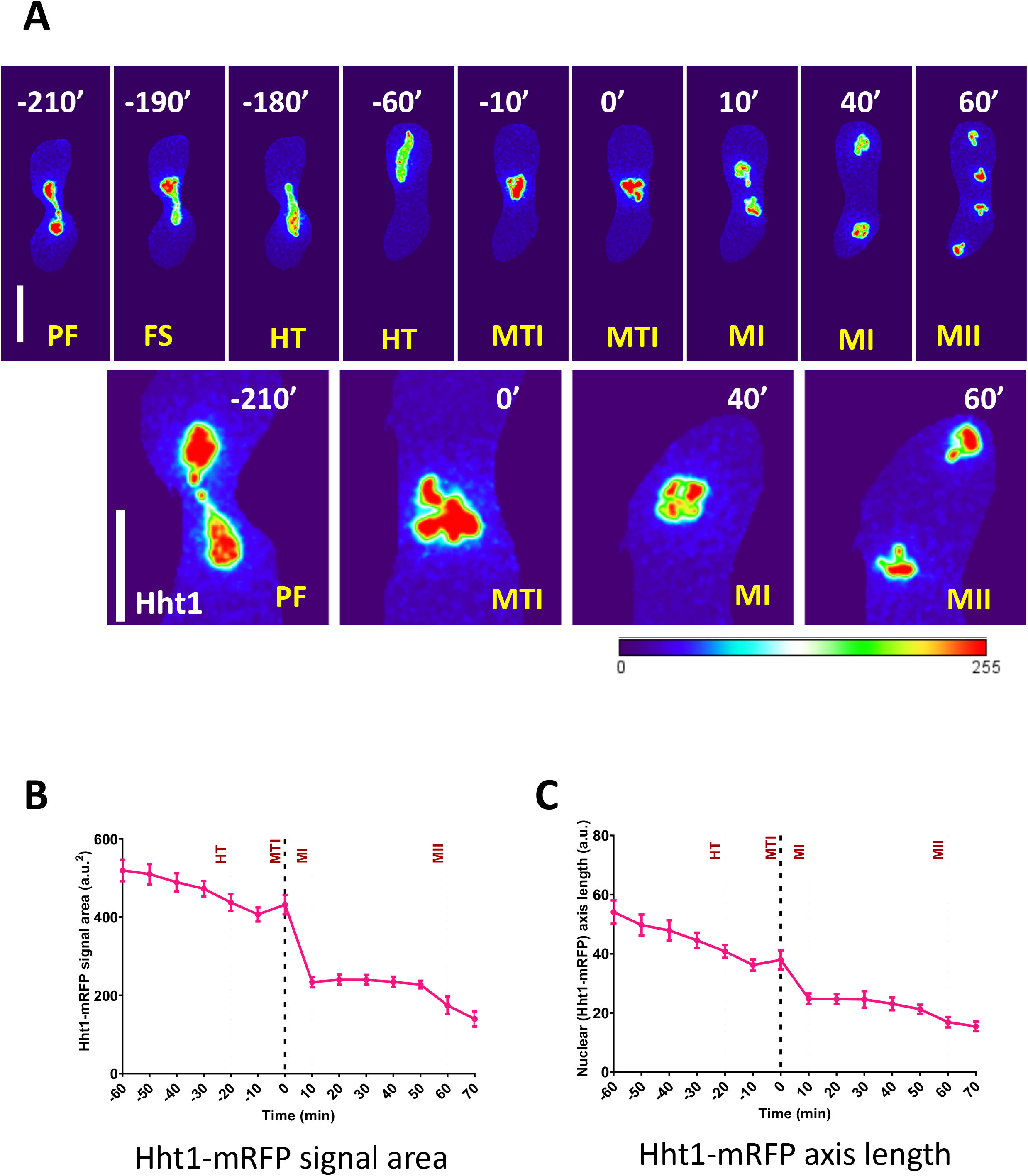
Nuclear dynamics of Hht1 during zygotic meiosis. (A) Panel showing selected time frames from Figure 1 (A) covering different meiotic signposts i.e. PF, FS, HT, MTI, MI and MII are chosen for optimal representation. Fiji’s “thermal” LUT was used to create false color images of defined timepoints which show a heatmap of fluorescence signal intensities for better understanding of nuclear dynamics. In the heatmap, red color is representing higher fluorescence signal intensity and highly compact nucleus. Second row is showing closeup panels of selected frames of row 1. The scale bars equal 5 μm. (B) Quantitation of Hht1-mRFP signal area. (C) The graph showing quantitation of nuclear size by using Feret’s diameter function of Fiji. Metaphase I is considered “t0” minutes. Timing prior ot MTI are denoted by negative signs while post MTI events are marked by Positive numbers. Values in the graphs represents mean and error bars represent 95% class intervals.

**Figure S2.**
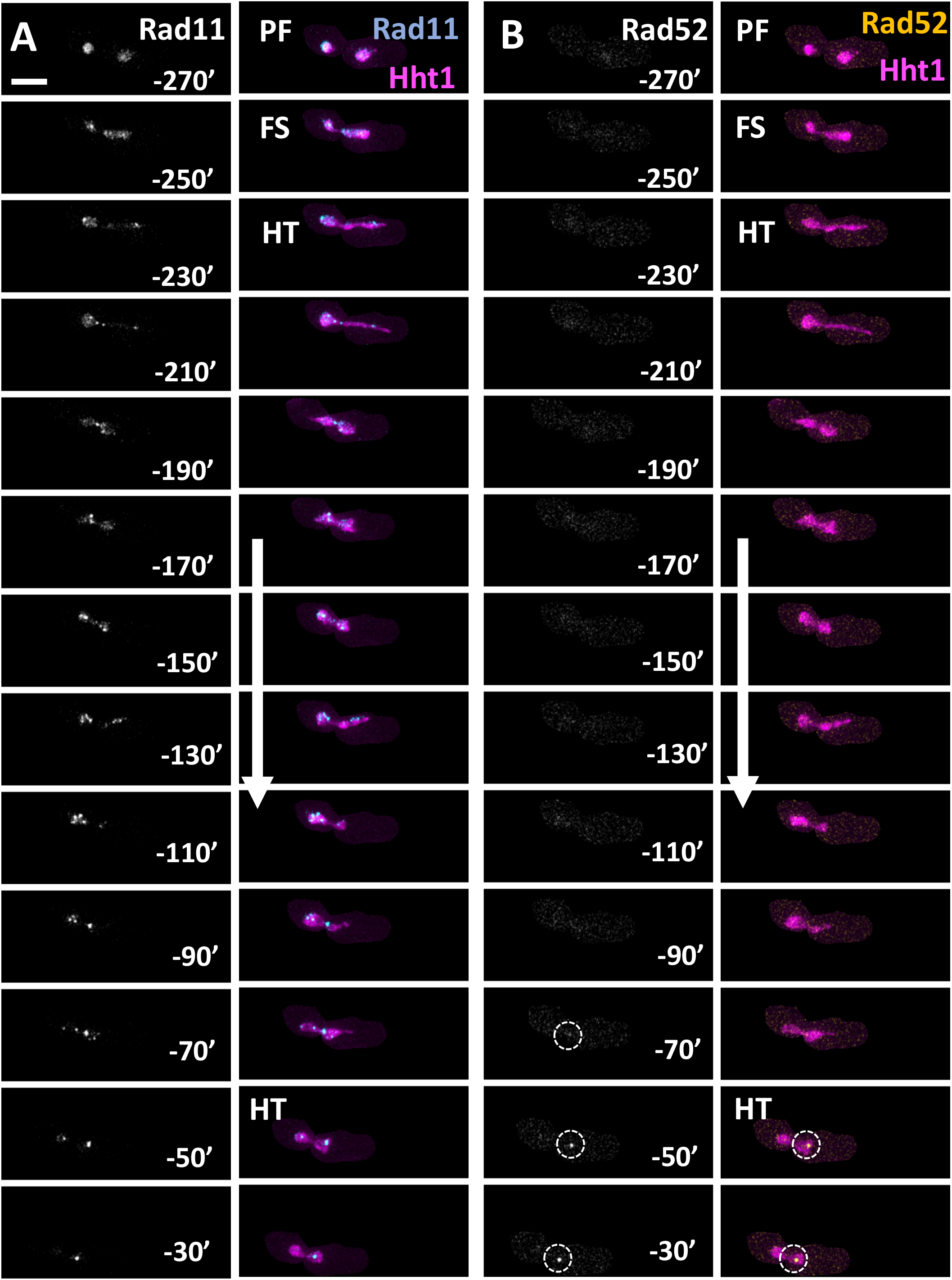
Dynamics of Rad11 and Rad52 foci. Panels showing time-lapse images of meiotic cells carrying fluorescently tagged histone (Hht1-mRFP), RPA (Rad11-CFP) and Rad52 (Rad52-YFP) (7327 × 7328). Panels (A) Rad11 and (B) Rad52 representing the dynamics of repair foci during different meiotic events encompassing prefusion (PF), fusion (FS), horsetail (HT) and metaphase I (MTI). Numbers represent timing in minutes and scale bars equal 5 μm.

**Figure S3.**
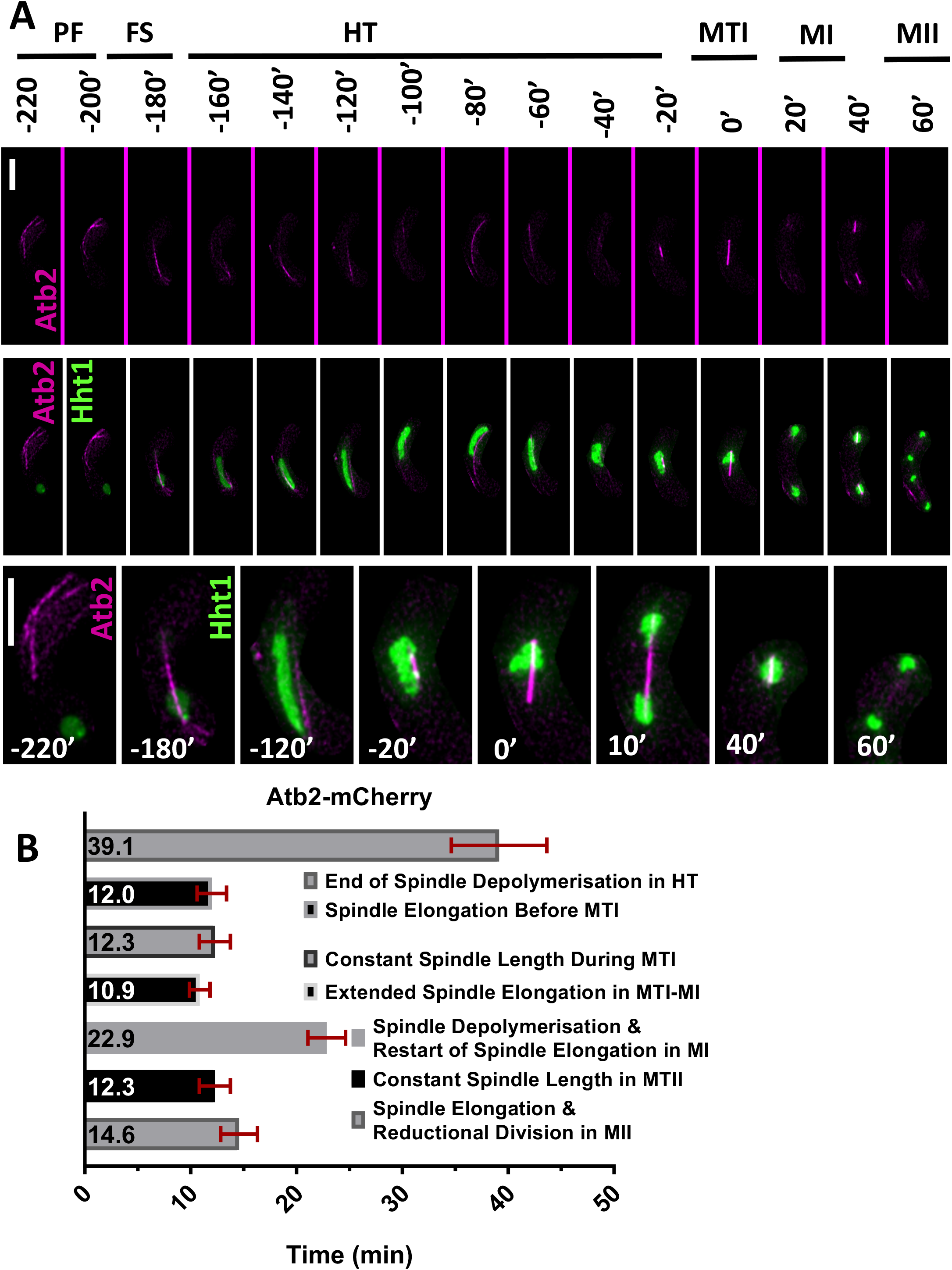
Microtubule dynamics. (A) Panels showing live-cell images of meiotic cells carrying fluorescently tagged histone (Hht1-GFP) and microtubule (Atb2-mCherry) (8055 × 8203). Time lapse images were captured every 10 minutes for 8 hours and selected frames are shown. Row 1 (Atb2-mCherry) and 2 (merged image of Atb2-mCherry Hht1-mRFP) show microtubule dynamics during different premeiotic and meiotic events covering pre-fusion to meiosis II (MII). Row 3 is showing the closeup images of selected time frames of row 2. Numbers represent timing in minutes and scale bars equal 5 μm. (B) Durations of microtubule depolymerization and elongation during different meiotic stages covering late horsetail (HT) to meiosis II (MII). More than 35 cells from at least two independent biological replicates were analyzed. Data represents mean timing of the microtubule dynamics. Error bars represent 95% class intervals.

**Figure S4.**
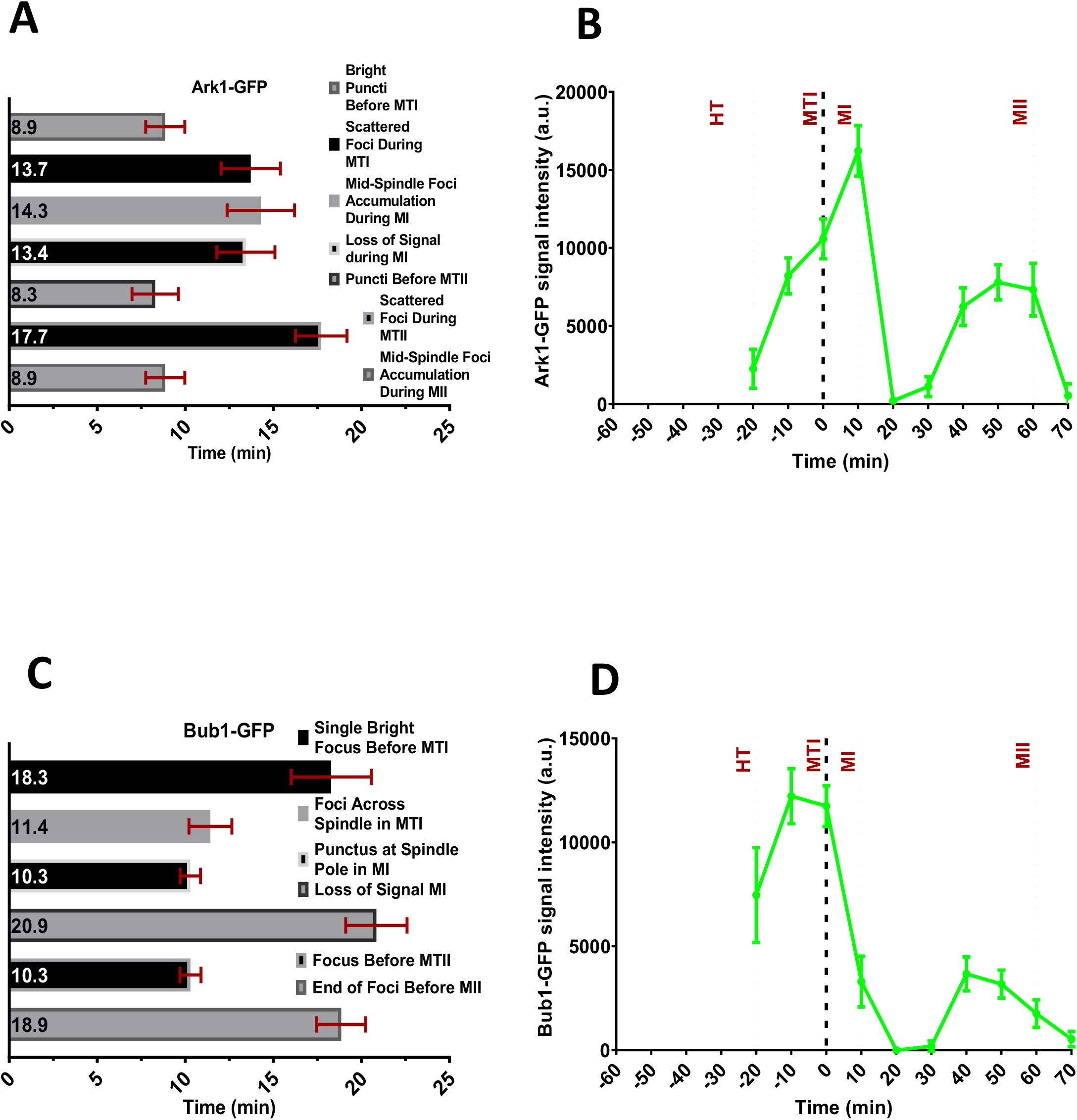
Aurora-B and Bub1 kinase dynamics. Panel (A) Ark1-GFP (8053 × 8056) and (C) Bub1-GFP (8055 × 8067) showing the duration of fluorescent signal dynamics during different meiotic events. Panel (B) Ark1-GFP and (D) Bub1-GFP showing the quantitation of fluorescent intensities. For both Ark1 and Bub1 dynamics, more than 35 cells of each from at least two independent movies were analyzed. Values presented in (A, C) and (B, D) are the means of timing and fluorescent intensities respectively. Error bars represent 95% class intervals.

**Table S1.**
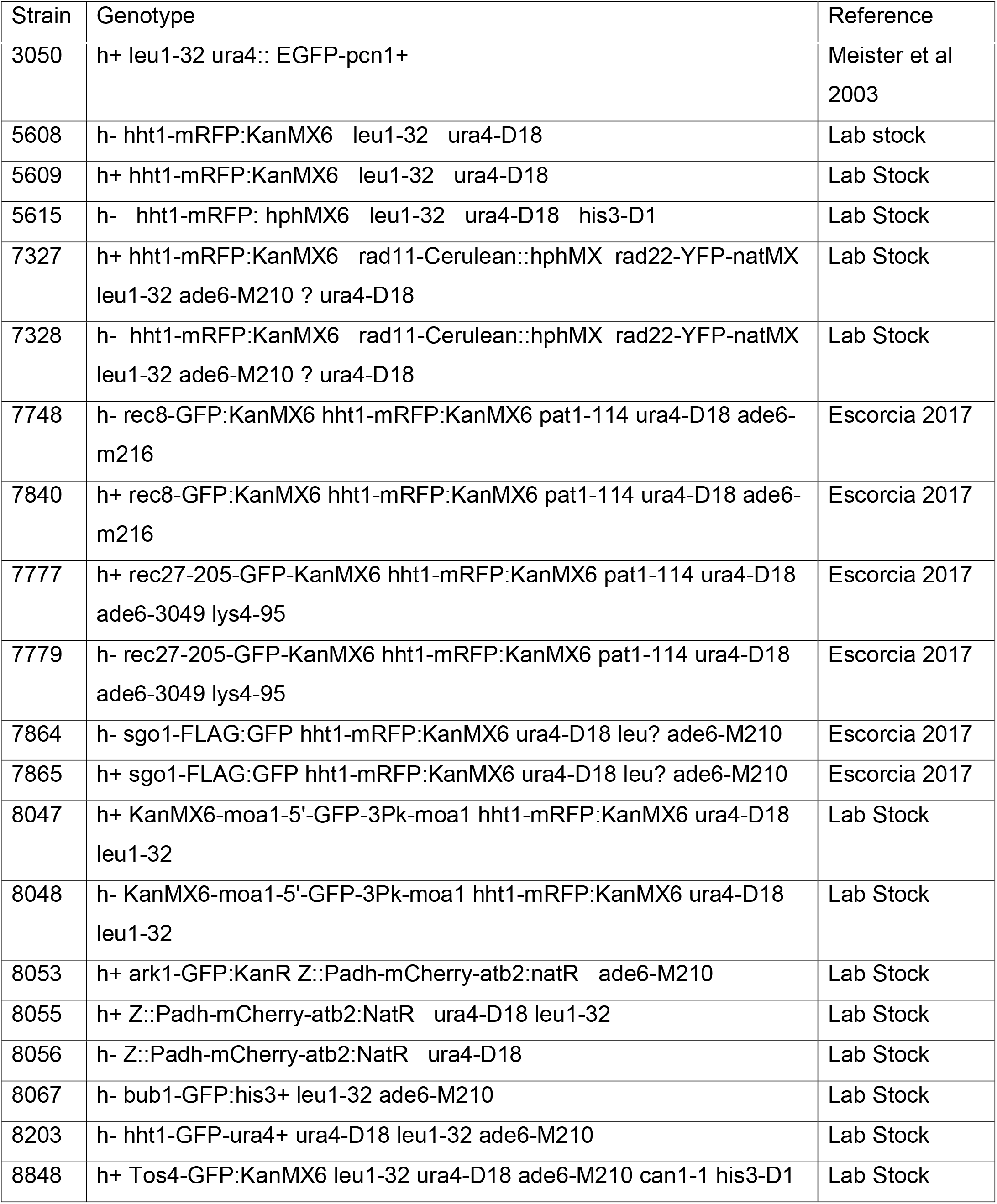
List of strains used in this study.

| Strain | Genotype | Reference |
| --- | --- | --- |
| 3050 | h+ leu1-32 ura4:: EGFP-pcn1+ | Meister et al<br>2003 |
| 5608 | h- hht1-mRFP:KanMX6 leu1-32 ura4-D18 | Lab stock |
| 5609 | h+ hht1-mRFP:KanMX6 leu1-32 ura4-D18 | Lab Stock |
| 5615 | h- hht1-mRFP: hphMX6 leu1-32 ura4-D18 his3-D1 | Lab Stock |
| 7327 | h+ hht1-mRFP:KanMX6 rad11-Cerulean::hphMX rad22-YFP-natMX<br>leu1-32 ade6-M210 ? ura4-D18 | Lab Stock |
| 7328 | h- hht1-mRFP:KanMX6 rad11-Cerulean::hphMX rad22-YFP-natMX<br>leu1-32 ade6-M210 ? ura4-D18 | Lab Stock |
| 7748 | h- rec8-GFP:KanMX6 hht1-mRFP:KanMX6 pat1-114 ura4-D18 ade6-<br>m216 | Escorcia 2017 |
| 7840 | h+ rec8-GFP:KanMX6 hht1-mRFP:KanMX6 pat1-114 ura4-D18 ade6-<br>m216 | Escorcia 2017 |
| 7777 | h+ rec27-205-GFP-KanMX6 hht1-mRFP:KanMX6 pat1-114 ura4-D18<br>ade6-3049 lys4-95 | Escorcia 2017 |
| 7779 | h- rec27-205-GFP-KanMX6 hht1-mRFP:KanMX6 pat1-114 ura4-D18<br>ade6-3049 lys4-95 | Escorcia 2017 |
| 7864 | h- sgo1-FLAG:GFP hht1-mRFP:KanMX6 ura4-D18 leu? ade6-M210 | Escorcia 2017 |
| 7865 | h+ sgo1-FLAG:GFP hht1-mRFP:KanMX6 ura4-D18 leu? ade6-M210 | Escorcia 2017 |
| 8047 | h+ KanMX6-moa1-5'-GFP-3Pk-moa1 hht1-mRFP:KanMX6 ura4-D18<br>leu1-32 | Lab Stock |
| 8048 | h- KanMX6-moa1-5'-GFP-3Pk-moa1 hht1-mRFP:KanMX6 ura4-D18<br>leu1-32 | Lab Stock |
| 8053 | h+ ark1-GFP:KanR Z::Padh-mCherry-atb2:natR ade6-M210 | Lab Stock |
| 8055 | h+ Z::Padh-mCherry-atb2:NatR ura4-D18 leu1-32 | Lab Stock |
| 8056 | h- Z::Padh-mCherry-atb2:NatR ura4-D18 | Lab Stock |
| 8067 | h- bub1-GFP:his3+ leu1-32 ade6-M210 | Lab Stock |
| 8203 | h- hht1-GFP-ura4+ ura4-D18 leu1-32 ade6-M210 | Lab Stock |
| 8848 | h+ Tos4-GFP:KanMX6 leu1-32 ura4-D18 ade6-M210 can1-1 his3-D1 | Lab Stock |

